# Nanophotonic neural probes for *in vivo* photostimulation, electrophysiology, and microfluidic delivery

**DOI:** 10.1101/2025.08.26.671930

**Authors:** Xin Mu, Homeira Moradi Chameh, Mandana Movahed, Fu Der Chen, John N. Straguzzi, Piyush Kumar, Andrei Stalmashonak, Hannes Wahn, Hongyao Chua, Xianshu Luo, Guo-Qiang Lo, Joyce K. S. Poon, Taufik A. Valiante, Wesley D. Sacher

## Abstract

Implantable silicon neural probes with integrated optical emitters and electrodes are emerging tools for simultaneous optogenetic stimulation and electrophysiological recording in deep brain regions. In parallel, neural probes with microfluidic channels have been developed for localized drug delivery and neurochemical sampling. However, thus far, such fluidic probes have lacked optical and electrical functionalities or been limited to a low number of optical emitters and/or electrodes, constraining their utility in multimodal investigations of neural circuits. Here, we introduce foundry-fabricated silicon nanophotonic neural probes with monolithically integrated microfluidics. Each probe has 16 silicon nitride grating coupler emitters, 18 titanium nitride microelectrodes, and one embedded microfluidic channel. We evaluate the photonic, electrophysiological, and microfluidic functionalities *in vivo* in optogenetic, blue-light-sensitive mice. With our multifunctional neural probes, we demonstrate local suppression of epileptic seizure activity (induced by microfluidic injection of 4-aminopyridine) using photostimulation. Through foundry-compatible microfluidics integration, this work advances the versatility of nanophotonic neural probes and presents new possibilities for multimodal neuroscience experiments leveraging this scalable neurotechnology.

## 1 Introduction

Deciphering brain function is a multiphysics challenge that requires interrogation of both superficial and deep structures. Optogenetics, enabling cell-type-specific, millisecond-scale optical control of neural activity [1, 2], combined with implantable neural probes for light delivery and electrical recording, provides powerful means for simultaneous modulation and monitoring of neuronal electrophysiology across brain regions [3–9]. However, a comprehensive understanding of brain function must also integrate neurochemistry, which shapes brain dynamics across physiological and pathological states and provides complementary routes for modulation of neural circuits. Recent advances in microchip-based silicon (Si) probes and fiber-based implants with integrated microfluidic channels have opened direct access to this domain [10–12]. Microfluidic delivery enables precise injection of chemicals and neuropharmacological agents into targeted brain regions, with different agents supporting diverse modes of neural circuit modulation, while also facilitating functional studies of pharmacological effects *in vivo* [13–15]. In parallel, microfluidic sampling quantifies neurochemicals essential for brain function (e.g. glutamate, *γ*-aminobutyric acid, and glucose) via microdialysis and chemical sensing [16–18], offering a path to link molecular signaling with circuit-level dynamics and neural responses. Integrating these capabilities with optical stimulation and electrophysiological recording into a single neurotechnology — *multimodal neural probes* — has emerged as a promising route toward unified optogenetic, neurochemical, and pharmacological interrogation, opening new avenues for neuroscience experiments.

While optical fibers with embedded microfluidic channels have demonstrated multimodal capabilities [12, 19], silicon neural probes offer a scalable platform that supports higher densities of emitters and microelectrodes within minimal implant volumes. These probes comprise narrow shanks (≈3–10 mm long, 30–100 µm thick, ≲100 µm wide) with arrays of optical emitters and electrodes, anchored to a base region that integrates the required circuitry and connection interfaces. Si neural probes enable precise, reconfigurable light delivery to deep brain regions, exceeding the tissue scattering limits of free-space photostimulation (one-photon: ∼100 µm; two-photon: ∼0.5–1 mm) [20]. They achieve implant volumes orders of magnitude smaller than gradient refractive index (GRIN) lenses [21] and comparable to tapered fibers [9]. Photostimulation is realized through integrated nanophotonic waveguides [3–5, 22, 23] or optoelectronic emitters (micro- or organic LEDs) [6–8] on neural probes, both intrinsically compatible with on-chip electrophysiological recording electrodes. Significant advances have been achieved in both approaches, with 256 optoelectronic emitters per shank reported in Ref. 7 and 28 nanophotonic waveguide emitters co-integrated with 960 electrodes on a single shank in Ref. 4. Yet, integrating microfluidics into these platforms while preserving state-of-the-art channel counts remains an open challenge. For example, in Ref. 13, a Si-based multimodal neural probe was reported with a single embedded microfluidic channel, 32 microelectrodes, and one polymer waveguide emitter. Other Si probes have integrated microfluidics with microelectrodes [15, 24]; however, photostimulation in Ref. 24 relied on a single co-packaged optical fiber, and no light-emission functionality was included in Ref. 15. Overcoming this challenge and realizing multimodal neural probes with dense, high-channel-count emitter arrays will unlock their full optical potential, adding spatially precise photostimulation of neuronal populations to their fluidic and electrophysiological recording functions.

In our prior work, we developed a series of nanophotonic neural probes of increasing complexity fabricated in a commercial Si photonics foundry. These neural probes guide light into the brain from external lasers using integrated photonic waveguides and diffractive (grating coupler) emitters. As the light sources (and their corresponding heat dissipation) are spatially separate from the shank and surrounding brain tissue, nanophotonic neural probes are capable of optical emission powers that surpass those of optoelectronic probes [5]. These higher powers provide flexibility to drive network-level activity and extend optical reach in scattering tissue [4, 25, 26]. For photonic routing, the probes integrate silicon nitride (SiN) waveguides, a standard Si photonic material transparent to both visible and near-infrared light [27–29]. Our earlier reports demonstrated the versatility of SiN grating coupler emitters for generating a variety of beam profiles: neuron-scale low-divergence beams [27], light sheets [30, 31], focused spots [32], and continuously-steerable beams [33]. The small dimensions and strong optical confinement of SiN nanophotonic waveguides naturally support dense photonic routing and compact emitters, enabling arrays with increasing density and channel count. Most recently, we introduced dual-color nanophotonic neural probes with 52 emitters (26 blue and 26 red), each delivering relatively high optical powers *>*80 µW per emitter, together with 26 recording microelectrodes co-integrated on a single shank [5]. This work complements parallel advances in nanophotonic neural probes, including implementations with on-chip optical switches [3], wavelength demultiplexing [34], and integration with CMOS-based neural recording [4], which offer additional strategies for scaling emitter and electrode channels.

Building on this foundation, we now present multimodal nanophotonic neural probes with photostimulation, electrophysiological recording, and microfluidic delivery functionalities, Fig. 1. Through monolithic integration of SiN waveguides, microelectrodes, and buried microfluidic channels (defined below dense on-shank waveguide routing and wiring), we achieve an emitter count surpassing previously reported multimodal neural probes. The neural probe chips are fabricated at wafer-scale at a commercial Si photonics foundry, enabling scalability in both complexity and manufacturing volumes. Each probe features 16 SiN grating coupler emitters, 18 microelectrodes, and one microfluidic channel integrated onto a single shank. The neural probe functionalities are demonstrated through *in vivo* experiments in optogenetic mice sensitive to blue-light photostimulation. Toward applications in epilepsy research, we induce seizure activity through microfluidic injection of an epileptogenic agent, achieve local seizure suppression via photostimulation, and monitor the process with electrophysiological recordings. Overall, this work introduces a foundry-compatible integration approach for incorporating microfluidics into advanced nanophotonic neural probes without compromising emitter or electrode densities, enhancing Si neural probe functionality and enabling new opportunities for multimodal neuroscientific investigations.

**Fig. 1:**
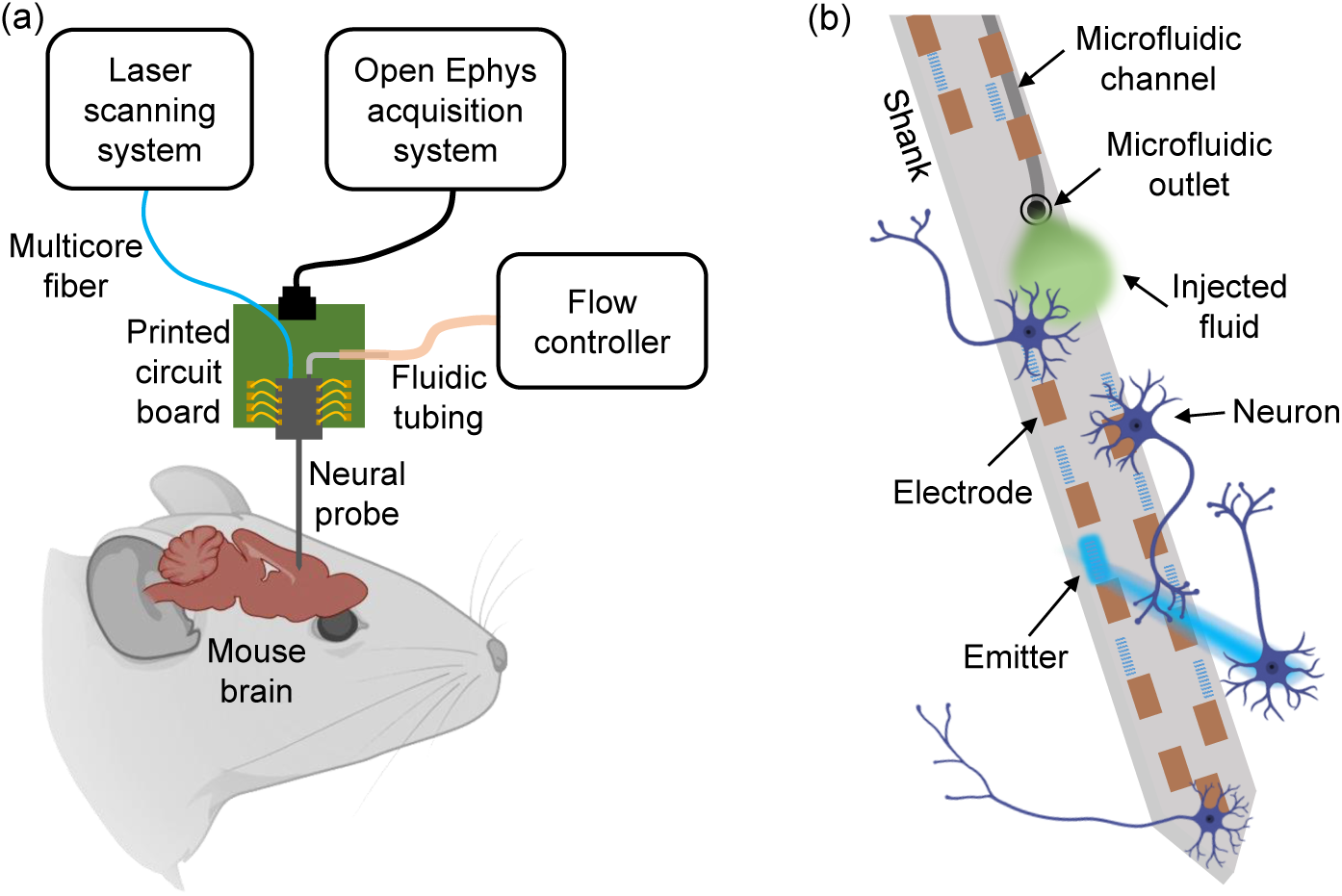
Conceptual illustrations of the multimodal neural probe. (a) *In vivo* experimental schematic with a packaged neural probe and the peripheral optical, electrophysiological recording, and fluidic subsystems. (b) Illustration of the neural probe operation. Beams are delivered from the optical emitters for targeted photostimulation, while fluid is injected via the embedded microfluidic channel. Microelectrodes distributed nearby the emitters and the microfluidic outlet detect changes in neural activity arising from the optical/chemical stimulation (not to scale). BioRender.com was used to generate the figures.

## 2 Results

### 2.1 Neural probe design, fabrication, and characterization

The neural probe chips were fabricated on 200-mm diameter Si wafers at Advanced Micro Foundry. The fabrication process is illustrated in Fig. 2a. Fabrication began with silicon dioxide (SiO_2_) deposition and formation of the microfluidic channels in the Si substrate through trench and isotropic etch steps. An additional SiO_2_ deposition step and chemical mechanical polishing (CMP) sealed the channels and formed the bottom cladding of the waveguides. Each sealed channel exhibited a cusp, which fused together within the SiO_2_ cladding, Fig. 2f. Next, two SiN waveguide layers (SiN1, 150-nm thickness; SiN2, 75-nm thickness; Fig. 2g) were fabricated using plasma-enhanced chemical vapor deposition, 193-nm deep-ultraviolet photolithography, and reactive ion etching. The SiO_2_ top cladding, three aluminum (Al) metal routing layers with corresponding vias, and titanium nitride (TiN) microelectrodes were formed subsequently. CMP was used for planarization of layers. A two-step etch of the SiO_2_ cladding opened microfluidic outlets. Lastly, the neural probe chip outlines were defined by deep trench etching, and the wafers were thinned to ≈100 µm by backgrinding to separate the chips. The deep trench etch step also exposed microfluidic channels at the facets of fluidic input ports, Fig. 2a. The two SiN waveguide layers were used for photonic routing, while three Al metal routing layers and microelectrodes (defined by TiN coating the topmost metal layer) were implemented for electrophysiological recording. Buried microfluidic channels beneath the photonic and electrical routing layers enabled fluid flow down the shank and into the brain via a microfluidic outlet (a trench locally opening the channel).

**Fig. 2:**
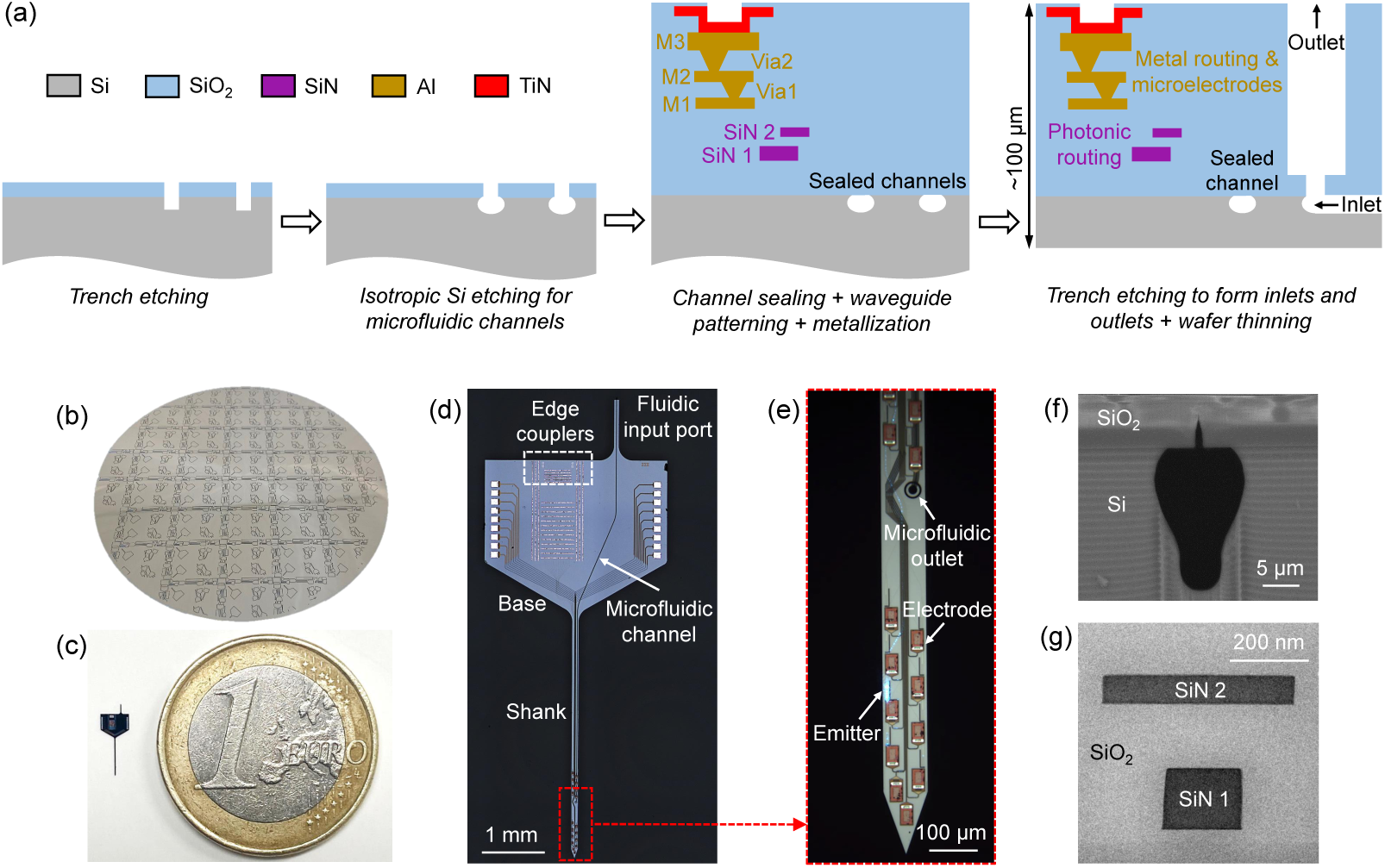
(a) Cross-section schematics of the neural probe fabrication process (not to scale). (b) Photograph of a 200-mm diameter neural probe wafer; the background was removed for improved visibility. (c) Photograph of a neural probe next to a coin. (d) Optical micrograph of a neural probe chip prior to packaging. (e) Enlarged view of the neural probe shank with one grating coupler emitting light. (f) Cross-section scanning electron micrograph of the integrated microfluidic channel. (g) Cross-section transmission electron micrograph of the two SiN waveguide layers. (d)-(g) adapted from our conference abstract, Ref. 35.

Each neural probe features 16 addressable nanophotonic grating coupler emitters, 18 TiN microelectrodes, and one microfluidic channel integrated onto a single 3.9-mm-long, 107-µm-wide shank, with corresponding photonic / electrical / fluidic routing defined on a 2.65 mm × 3 mm base region. A 1-mm-long, 80-µm-wide fluidic input port extends from the base of the probe, enabling coupling to external fluidic tubing. The buried microfluidic channel (averaged cross-sectional area ≈160 µm^2^) traverses the fluidic input port, base region, and shank — terminating with a 16-µm-diameter outlet on the shank, Fig. 2e. An array of 16 fiber-to-chip, bi-layer, waveguide edge couplers (SiN1 and SiN2 layers) [36] along the probe base facet couple laser light onto the neural probe chip from a custom 16-channel multicore optical fiber [37] (each fiber core being aligned to a corresponding edge coupler). Following the edge couplers, each waveguide path includes a polarizer consisting of 44 cascaded SiN1 90° bends (18-µm radius, 300-nm waveguide width) for preferential transmission of transverse-electric (TE) polarized light. SiN1 routing waveguides direct light from the base region to grating coupler emitters on the shank. To reduce optical scattering loss and inter-waveguide crosstalk, the routing waveguides were tapered to dissimilar interlaced widths of 600 and 700 nm. Each grating coupler emitter (6 µm wide and 25 µm long) was designed with a 440-nm grating pitch and 50% duty cycle, forming a low-divergence emitted beam [27]. The grating emitters are staggered on a 55-µm longitudinal inter-grating pitch along the shank. 16 TiN electrodes (26.7 µm × 15 µm) for electrophysiological recording are distributed in the vicinity of the 16 grating emitters, with 2 additional TiN electrodes located near the shank tip. The microfluidic outlet is positioned in the middle of the array of emitters and electrodes (≈0.9 mm from the shank tip), with the aim of maximizing the overlap of the diffusion profile of injected fluid with the array (spanning 1.35 mm along the shank).

A fully packaged neural probe mounted on a printed circuit board (PCB) with an attached fluidic tube and multicore optical fiber is shown in Fig. 3a. The packaging process is detailed in Section 4.2. A schematic of the connections between the neural probe and the peripheral optical, electrophysiological recording, and fluidic subsystems is shown in Fig. S1, with accompanying details in Sections 4.3 and 4.4. Briefly, electrophysiological signals are routed from the on-chip recording electrodes to a data acquisition system (Open Ephys [38]) via the PCB, a recording headstage circuit board (for amplification and digitization), and an electrical cable. The fluidic tubing is connected to a flow controller for fluid delivery. Concurrently, laser light from a 488-nm-wavelength diode laser is coupled to the neural probe chip through a computer-controlled laser scanning system and multicore fiber; each fiber core aligned and coupled to an on-chip edge coupler of the neural probe chip. The scanning system selects the active core of the fiber, routing light to the corresponding on-chip waveguide and grating coupler emitter (addressing the emitters on the neural probe). Additional components in the optical subsystem (shutter, attenutator, and depolarizer) define the photostimulation patterns and emission powers, while circumventing polarization fluctuations in the multicore fiber.

**Fig. 3:**
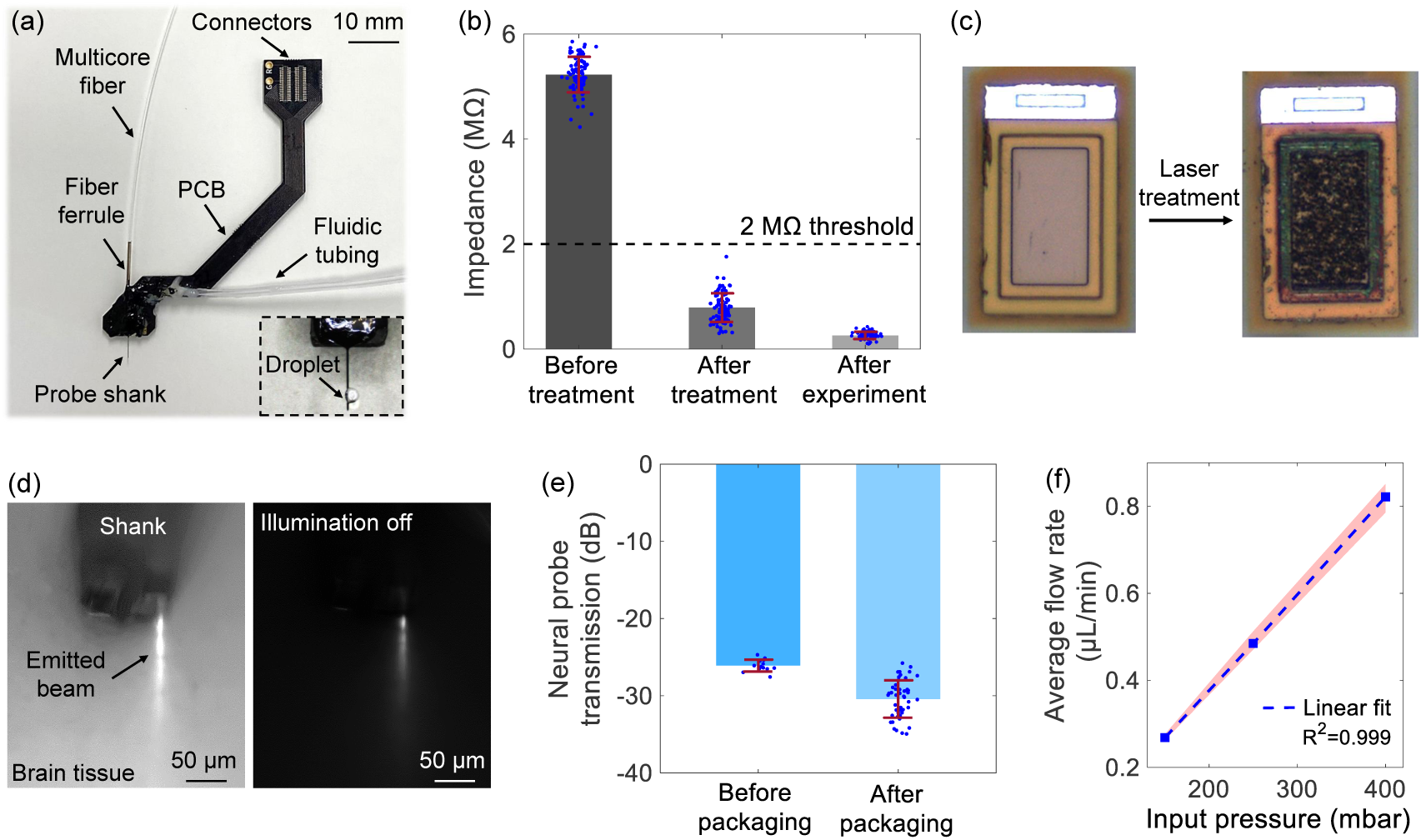
Neural probe characterization. (a) Photograph of a packaged neural probe. (Inset) A droplet of injected fluid near the outlet of a second packaged probe. (b) Electrochemical impedance of the electrodes before laser treatment, after laser treatment, and after *in vivo* experiments. Before treatment: n=80 electrodes from 6 probe samples; after treatment/experiment: n=92 electrodes from 6 probes. (c) Micrographs of electrode surfaces before and after laser treatment; adapted from Ref. 35. (d) Top-down fluorescence microscope images of an emitted beam in a brain slice from a Thy1-ChR2-EYFP mouse with additional microscope epi-illumination on (left) and off (right). (e) Optical transmission of neural probes before and after packaging. Before packaging: emission power measured with TE-polarized light; n=13 emitters from 5 probe samples. After packaging: emission power measured with a depolarized laser input; n=48 emitters from 3 probes. (f) Flow rate vs. input pressure during *in vivo* microfluidic injection (from 3 probes). The error bars represent the standard deviations (SDs).

Prior to neural probe packaging, electrodes on the neural probe were post-processed with femtosecond laser pulses to achieve roughened electrode surfaces with low electrochemical impedance (see Section 4.1) [5, 26]. The as-fabricated TiN electrodes had impedances of 5.23 ± 0.34 MΩ at a frequency of 1 kHz (Fig. 3b), which was higher than the 2-MΩ threshold for electrophysiological recording with high signal-to-noise ratios [39]. Electrode impedances were reduced to 0.79 ± 0.27 MΩ after the laser treatment, and further reduced to 0.26 ± 0.07 MΩ after *in vivo* experiments (described in Sections 2.2 - 2.4). Reduced electrode impedance after *in vivo* experiments was also observed in our previous work [26], and we attribute this to rinsing of the neural probes in a cleaning solution (Tergazyme) after each experiment. Micrographs of a TiN electrode surface before and after laser treatment are shown in Fig. 3c.

Profiles of emitted beams from a packaged neural probe were measured in brain slices from a 4-month-old transgenic mouse expressing Channelrhodopsin-2 in pyramidal cells with a yellow fluorescent protein labeling (Thy1-ChR2-EYFP mouse), Fig. 3d. Details of the beam profile measurements are listed in Section 4.6 and in our previous work [26]. The averaged full width at half maximum of the emitted 488-nm-wavelength beams at a 100-µm propagation distance in cortex were ≈55 µm. The narrow beam widths are approximately matched to the emitter pitch on the shank and enable optogenetic stimulation with few-neuron spatial resolution and high fill factor. The optical transmission of neural probe chips was measured before and after packaging (Fig. 3e). We define transmission as the ratio of output optical power from a grating coupler emitter to the input optical power at the edge coupler facet; measurements were performed at a wavelength of 488 nm. The optical transmission was −26.1 ± 0.8 dB (mean ± standard deviation) for grating emitters from bare neural probe chips before packaging with TE-polarized light. After packaging, the optical transmission was measured as −30.4 ± 2.4 dB across grating emitters with a depolarized optical input (to mimic the conditions of the *in vivo* experiments). ≈10 dB of the optical loss is attributed to the intrinsic loss of the fiber-to-chip couplers; on-chip waveguide routing, polarizers, and grating emitters further add to the loss. Following packaging, the neural probes exhibited higher losses and higher transmission variation, resulting from misalignment between the multicore fiber and the edge couplers during and after the fiber attachment process.

Figure 3f shows the measured flow rate versus input pressure during microfluidic injection into brain tissue *in vivo*. The high linearity between flow rates and pressure are indicative of the absence of leakage or clogging in the microfluidic channels. The averaged flow resistance of the microfluidic channels (≈450 mbar*min/µL) is moderate compared with other neural probes with microfluidic functionalities [12, 13, 19, 24], consistent with the relatively moderate dimensions of the channel cross-section (see Table 1).

**Table 1:** Comparision of multimodal implantable Si- and fiber-based neural probes with light emission, electrophysiological recording, and microfluidic functionalities.

| Device design | Implant cross-sectional area | No. of electrodes | No. of emitters | No. of microfluidic channels | Cross-sectional area of the channel | Ref. |
| --- | --- | --- | --- | --- | --- | --- |
| Four-shank Si probe | 128 $\mu\text{m} \times 40 \mu\text{m}$<br>$\times 4$ shanks | 32<br>(8/shank) | 1 polymer waveguide | 1 multi-cavity channel | 10 $\mu\text{m} \times 12 \mu\text{m}$<br>for each cavity | [13] |
| Assembly of a glass fiber and a Si probe | 150 $\mu\text{m} \times 50 \mu\text{m}$<br>+ 225-$\mu\text{m}$ diameter fiber | 32 | 1 fiber core | 1 | 11 $\mu\text{m} \times <5 \mu\text{m}$ | [24] |
| Polymer fiber neural probe | $\approx 160 \mu\text{m}$ diameter | 4 | 8 fiber cores | 1 | $\approx 80\text{-}\mu\text{m}$ diameter | [19] |
| Polymer fiber neural probe | 306 $\mu\text{m} \times 342 \mu\text{m}$ | 6 | 1 fiber core | 1 | 100 $\mu\text{m} \times 40 \mu\text{m}$ | [12] |
| Si nanophotonic neural probe | 107 $\mu\text{m} \times 100 \mu\text{m}$ | 18 | 16 SiN grating coupler emitters | 1 | $\approx 160 \mu\text{m}^2$ | This work |

### 2.2 Optogenetic stimulation and electrophysiological recording

To firstly verify the optogenetic stimulation and electrophysiological recording functionalities of our neural probes, we conducted *in vivo* experiments in awake, head-fixed Thy1-ChR2-EYFP mice (2–4 months old, male and female). This optogenetic transgenic line broadly expresses the opsin Channelrhodopsin-2 (ChR2) in excitatory pyramidal neurons across the cortex and is widely used in neuroscience [40–42], providing a well-established model in which photostimulation evokes robust, easily detectable increases in neural activity, in contrast to inhibitory effects that are less readily detectable. The neural probes were mounted on a 4-axis micromanipulator and implanted between the somatosensory and motor cortex [anterior–posterior (AP): −0.5 mm, medial–lateral (ML): 1.2 mm] to depths of ≈ 1.0 - 1.9 mm, as measured from the shank tip. The details of the experimental procedure are described in Section 4.5; the generation of photostimulation patterns and recording of electrophysiological signals are detailed in Section 4.3.

Validation of photostimulation and electrophysiological recording capabilities of the neural probes was performed with 5 mice *in vivo*. Figure 4 shows representative results from one of the experiments. The photostimulation pattern consisted of 10 optical pulses at a 5 Hz frequency. The pulse width was 30 ms and the recovery time between two pulse trains was 10 s. The pattern was repeated 15 times for each selected optical emitter in Fig. 4a (Emitters A to D). Across the 60 photostimulation patterns applied, only one emitter was addressed during each repeat of the pattern, and the sequence of emitter selection was randomized. The recorded electrophysiological voltage traces were processed with common average referencing, bandpass filtering (300 - 6000 Hz), and photostimulation artifact removal. Processed electrophysiological recording data underwent spike sorting and analysis (see Section 4.7 for data analysis procedure). Waveforms and autocorrelograms of two example sorted single units (Unit 1 and Unit 2) are presented in Figs. 4b and 4c, with a low interspike interval violation ≤ 0.5 [43]. Unit 1 was recorded on Electrode 6 (closest to Emitter C), while Unit 2 was captured by Electrode 9 (near Emitter D). Figure 4d shows the spike raster plot of Unit 1 with 150 optical pulses (10 pulses × 15 repeats) from Emitter C. 91% of spikes recorded in a 230 ms acquisition window occured during the 30 ms optical pulse, showing robust activation of spiking events during the optical pulse.

**Fig. 4:**
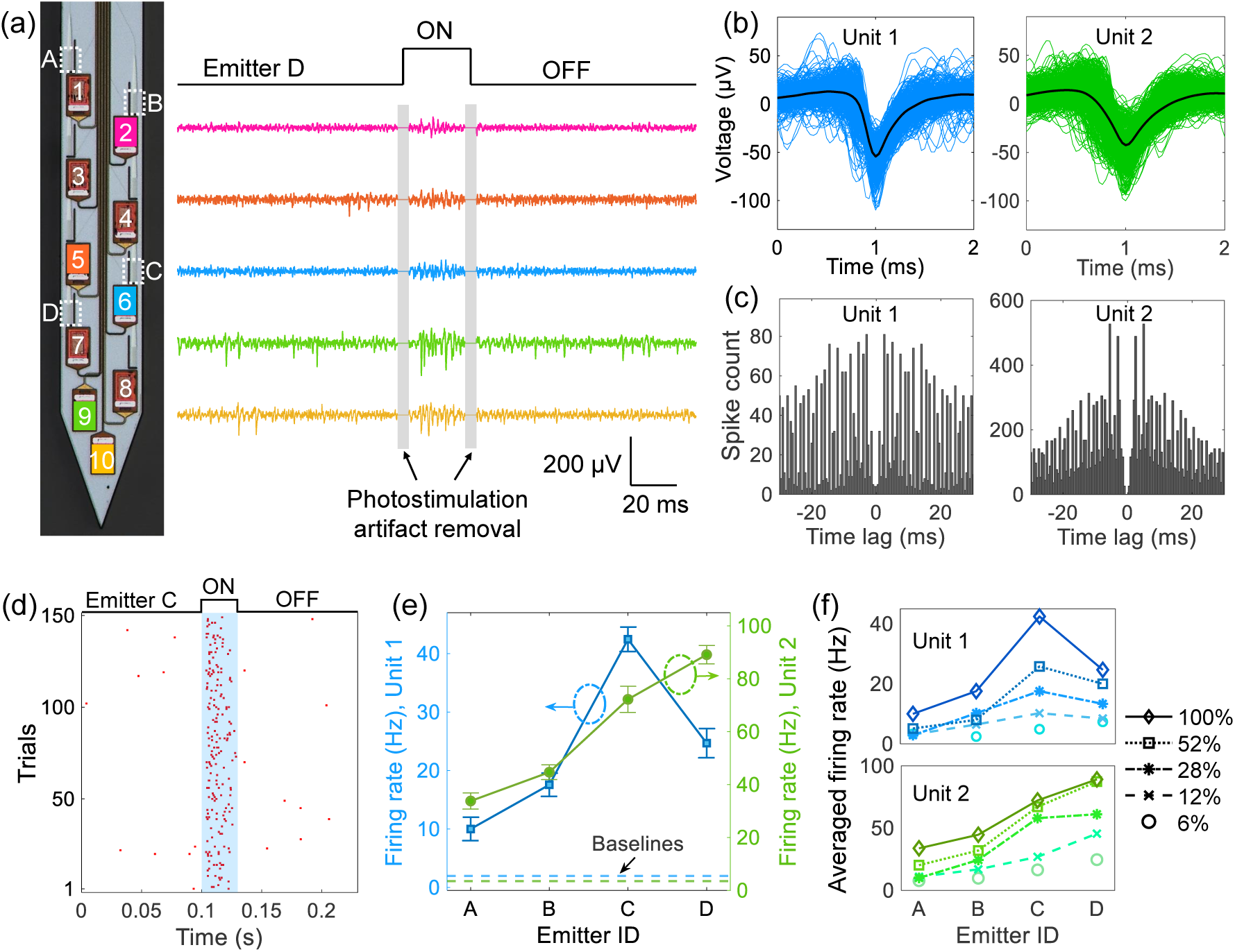
*In vivo* photostimulation and electrophysiologcal recording validation. (a) (Left) Micrograph of probe shank with selected electrodes (1 - 10) and emitters (A - D) labeled. (Right) Voltage traces from 5 selected electrodes showing evoked spikes during a 30-ms optical pulse from Emitter D at 28% optical power (emission power ≈ 1.3 µW). The displayed voltage traces were processed with common average referencing, bandpass filtering (300 - 6000 Hz), and photostimulation artifact removal. (b) Waveforms of two sorted single units (Units 1 and 2) from electrodes 6 and 9, respectively. (c) Autocorrelograms of the two sorted units. (d) Spiking raster plot of Unit 1 during the 30 ms photostimulation period over 150 pulses with Emitter C at 100% optical power (≈ 4.9 µW). (e) Mean firing rate of the two sorted units with photostimulation from different emitters at 100% optical power. Mean baseline firing rates are averaged over 10 s before each stimulus. Error bars represent the standard error of the mean (SEM). (f) Power dependence of mean firing rates of Units 1 and 2 (averaged over 150 optical pulses from Emitters C and D, respectively).

Figure 4e shows mean firing rates of the two sorted units under photostimulation from the 4 selected emitters labeled in Fig. 4a. Stimulated firing rates were higher than the spontaneous activity baselines for both units, and the change in firing rate increased with proximity of the emitter to the unit, implying spatial selectivity of photostimulation via addressing of emitters on the neural probe. In addition, the dependence of firing rates on photostimulation power was investigated (Fig. 4f) with 5 successively higher optical power levels (listed as a percentage of the maximum power with Emitter C, ≈4.9 µW). Optical power was controlled with the variable optical attenuator in Fig. S1. Both sorted units showed increased firing rates with higher optical power and the largest changes occured for the emitter closest to the unit.

Photostimulation with spatial selectivity and power dependence was also observed during the other 4 *in vivo* experiments. For sorted single units among all 5 experiments, the firing rate increased by a factor of ≈ 3.6 - 20.0 as the emission power of the closest emitter was increased from the lowest to highest levels tested. Control experiments were performed using the same photostimulation pattern (at similar optical power levels) in awake head-fixed wild-type mice. Photostimulated neural activity was not observed in the control experiments (see Fig. S2), indicating that the recorded neural activation with light delivery stemmed from optogenetic (rather than thermal) stimulation. In summary, the experiments in this section validate the electrophysiological recording and spatially-selective photostimulation functionalities of our nanophotonic neural probes. While these capabilities enable addressable optical modulation and monitoring of neural activity, targeted delivery of pharmacological agents provides a complementary route for modulating neural circuits. We therefore next report *in vivo* validation of the probes’ microfluidic injection functionality.

### 2.3 Microfluidic injection of 4-AP for seizure induction

Implantable neural probes with integrated microfluidic channels enable precise spatial and quantitative delivery of neuropharmacological agents and other chemicals into targeted brain regions, providing new opportunities to study neural activity under chemical modulation [10]. Epilepsy, which affects over 60 million people worldwide [44], is a significant neurological disorder whose impact underscores the need for continued research into its mechanisms. Epileptic seizures, the defining symptom of epilepsy, are characterized by abnormal, synchronized patterns of brain activity [45]. In animal models, seizures can be induced by local drug administration and studied through concurrent electrophysiological recording, providing a controlled approach to study their initiation and progression [10]. 4-aminopyridine (4-AP), a voltage-gated K^+^ channel blocker, is one of the most widely used seizure-inducing agents [46]. In this section, we validate the microfluidic functionality of our multifunctional neural probe and highlight its relevance to epilepsy research by inducing seizures *in vivo* through targeted microfluidic injection of 4-AP.

12 experiments were performed with awake head-fixed Thy1-ChR2-EYFP mice (2 - 4 months old, male and female). For each *in vivo* experiment, a solution of 15 mM 4-AP and 25 µM Alexa Fluor 568 conjugated dextran fluorescent dye in 1× phosphate buffered saline (PBS) was loaded into the microfluidic channel of the neural probe through the flow control system, Fig. 5a. As in Section 2.2, the neural probe was implanted between the somatosensory and motor cortex (AP: −0.5 mm, ML: 1.2 mm) of the mouse brain (Figs. 5b and 5c). The solution of 4-AP and fluorescent dye was injected into the brain at flow rates ranging from 0.2 - 1.0 µL/min under low pressures of 100 - 400 mbar, limiting tissue displacement and damage during the injections. Chemically induced seizures in cortical layers were successfully demonstrated in all 12 experiments, and Fig. 5 presents results from a representative experiment.

**Fig. 5:**
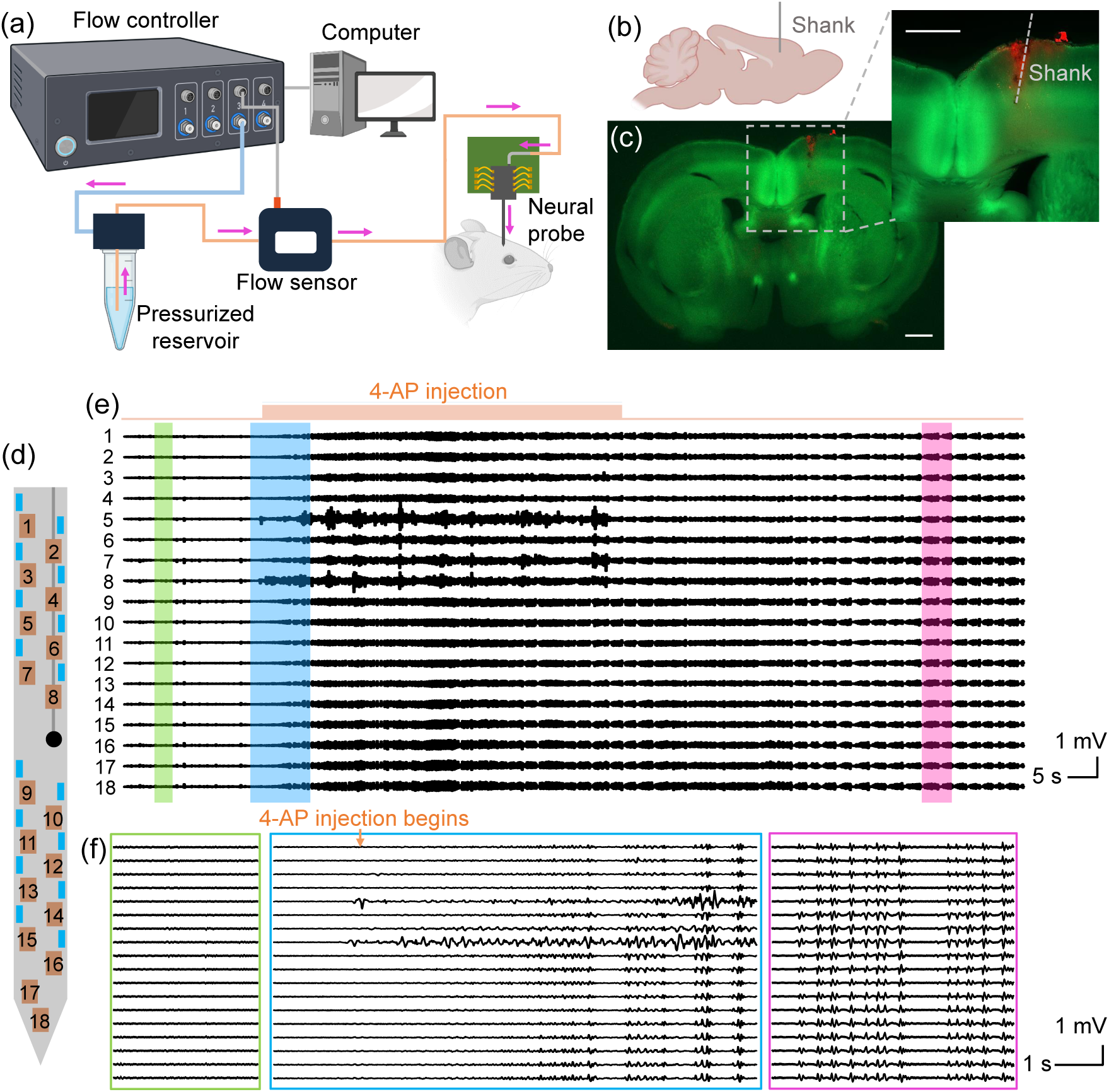
*In vivo* microfluidic injection experiment. (a) Schematic of the flow control system. (b) Conceptual image showing the implant location of the neural probe (sagittal plane). (c) Fluorescence images of a coronal brain slice including the implant site showing YFP fluorescence (green) and injected dye fluorescence (red). Scale bars: 1 mm. (d) Probe shank schematic showing the microfluidic outlet (black circle) and labeled electrodes (1 - 18). (e) Local field potential (LFP) traces from 18 electrodes following a 1-min 4-aminopyridine (4-AP) injection, showing seizure induction with microfluidic injection. The displayed voltage traces are processed with common average referencing and bandpass filtering (5 - 300 Hz). (f) Snapshots of the LFP traces before 4-AP injection (resting stage, green), upon 4-AP injection (blue), and during the induced seizure (epileptiform activity, magenta). BioRender.com was used to generate panels (a) and (b).

Figure 5a shows a schematic of the flow control system for microfluidic injection with the neural probe (additional details in Section 4.4). The input pressure was detected by the pressure-driven flow controller and flow rates were acquired with an in-line flow rate sensor in real time. The feedback control loop between the flow controller, laboratory computer, and flow sensor ensured stable and precise flow initiation, infusion, and termination. A disposable 1.5-ml Eppendorf tube was used as a fluid reservoir. During the experiment in Fig. 5, the fluidic injections were repeated with an averaged flow rate of 0.31 µL/min (1 min per injection, 5-min interval) until seizure activity was observed in electrophysiological recordings. Figure 5e shows the local field potential (LFP) traces recorded from 18 electrodes on the neural probe, following processing with common average referencing and 5 - 300 Hz bandpass filtering. During the second injection (at a total injected 4-AP solution volume of ≈0.61 µL), seizure activity was observed in the LFP recordings. At seizure onset, the LFP pattern transitioned to synchronized bursting activity across multiple recording electrodes, characterized by high-amplitude spikes and rhythmic after-discharges — indicative of successful seizure induction. Seizure behaviors in the mouse were also observed with forelimb clonus, corresponding to Racine Scale 3 behaviors [47] in rodent models of epilepsy. Figure 5f shows LFP signals at the resting stage, at seizure onset, and during epileptiform activities, respectively. LFP signals from electrodes 5 to 8 (close to the microfluidic outlet) exhibited larger amplitudes during the 4-AP injection, which may result from recording artifacts during fluid injection. Successful seizure induction via microfluidic 4-AP injection was also observed in the other 11 *in vivo* experiments, and seizures initiated after 1 - 4 injections and a total 4-AP injection volume ranging from ≈ 0.23 - 2.7 µL. The variation in the total number and volume of 4-AP injections required to induce seizures may be due to variations in the body weight and gender of the mice [48]. With seizure induction, epileptiform activities were observed on all electrodes within the brain (varying from 13 to 18, depending on the implantation depth).

Following the experiment, the mouse brain was extracted and 300-µm-thick coronal slices were prepared. Fluorescence imaging was performed to locate the neural probe insertion tract and confirm the microfluidic injection. An overlaid fluorescence image with yellow fluorescent protein (YFP) fluorescence from the brain (co-expressed with Channelrhodopsin-2 in the transgenic mice) and red fluorescence of the injected dye is shown in Fig. 5c, verifying the implant coordinate and successful fluid delivery. A solution of 25 µM Alexa Fluor 568 conjugated dextran dye was injected at similar flow rates and doses in control experiments with Thy1-ChR2-EYFP mice. No distinct changes in electrical patterns were observed during the experiments, indicating the seizure induction originated from the effects of 4-AP rather than the dye, PBS, and/or mechanical pertubration from the injection. Overall, this experiment validates the ability of our neural probes to induce seizures in targeted brain regions via drug delivery while simultaneously recording electrophysiological responses, establishing their microfluidic injection functionality and relevance to epilepsy research.

### 2.4 Local suppression of 4-AP-induced seizure with continuous-wave photostimulation

In the previous sections, we demonstrated the individual functionalities of our neural probes for spatially-selective optogenetic stimulation and seizure induction via targeted microfluidic injection. Building on this foundation, we next report *in vivo* experiments demonstrating their full multimodal operation, including seizure induction with 4-AP, photostimulation for local seizure suppression, and electrophysiological recording of chemically-induced and optically-modulated activity. This work further serves as an example application of the neural probes aligned with recent neuroscientific research studying seizure suppression in mouse models of epilepsy; via drug delivery [46], electrical stimulation [49, 50], and optogenetic stimulation [50–52]. Suppression of 4-AP-induced seizures with relatively long, “continuous-wave” (CW), optical pulses emitted from the neural probe was demonstrated in 5 experiments with awake head-fixed Thy1-ChR2-EYFP mice (2 - 4 months old, male and female).

Figure 6 presents results from a representative experiment. The target implant coordinate was identical to Sections 2.2 and 2.3, and the 4-AP and fluorescent dye solution of Section 2.3 was used for the microfluidic injections. Multiple injections were conducted with increasing flow rates from 0.28 - 1.0 µL/min (1-min injection times at a 5-min interval). Mouse behavior and local field potential were monitored to identify the onset of seizure activity, which was observed after the fourth injection (with a total injected solution volume of ≈2.7 µL). When sustained after-discharges were confirmed, photostimulation from the neural probe was applied for seizure suppression. The photostimulation pattern consisted of 10-s CW pulses delivered from individual emitters sequentially. 6 selected emitters were addressed in a randomized sequence, Fig. 6a, and the recovery time was 30 s between optical pulses. The photostimulation pattern was repeated at least 3 times during each experiment and emission powers ranged from ≈ 2.4 - 6.4 µW. Figures 6a and 6b show example processed LFP signal traces recorded from 17 electrodes (during seizure activity) before, during, and after a 10-s CW optical pulse from a single emitter. Generally, the amplitudes of the LFP signals during photostimulation were lower compared to pre- and post-stimulation, indicating local suppression of seizure activity with light delivery. It is noteworthy that the LFP signals in Fig. 6b exhibit a polarity inversion between the upper and lower electrode channels, with the transition centered around the microfluidic outlet. This pattern is consistent with sink-source relationships across the electrode channels with possible dependencies on the formation of an epileptic focus in the vicinity of the microfluidic outlet [53, 54] in addition to the microelectrodes spanning multiple cortical layers [55].

**Fig. 6:**
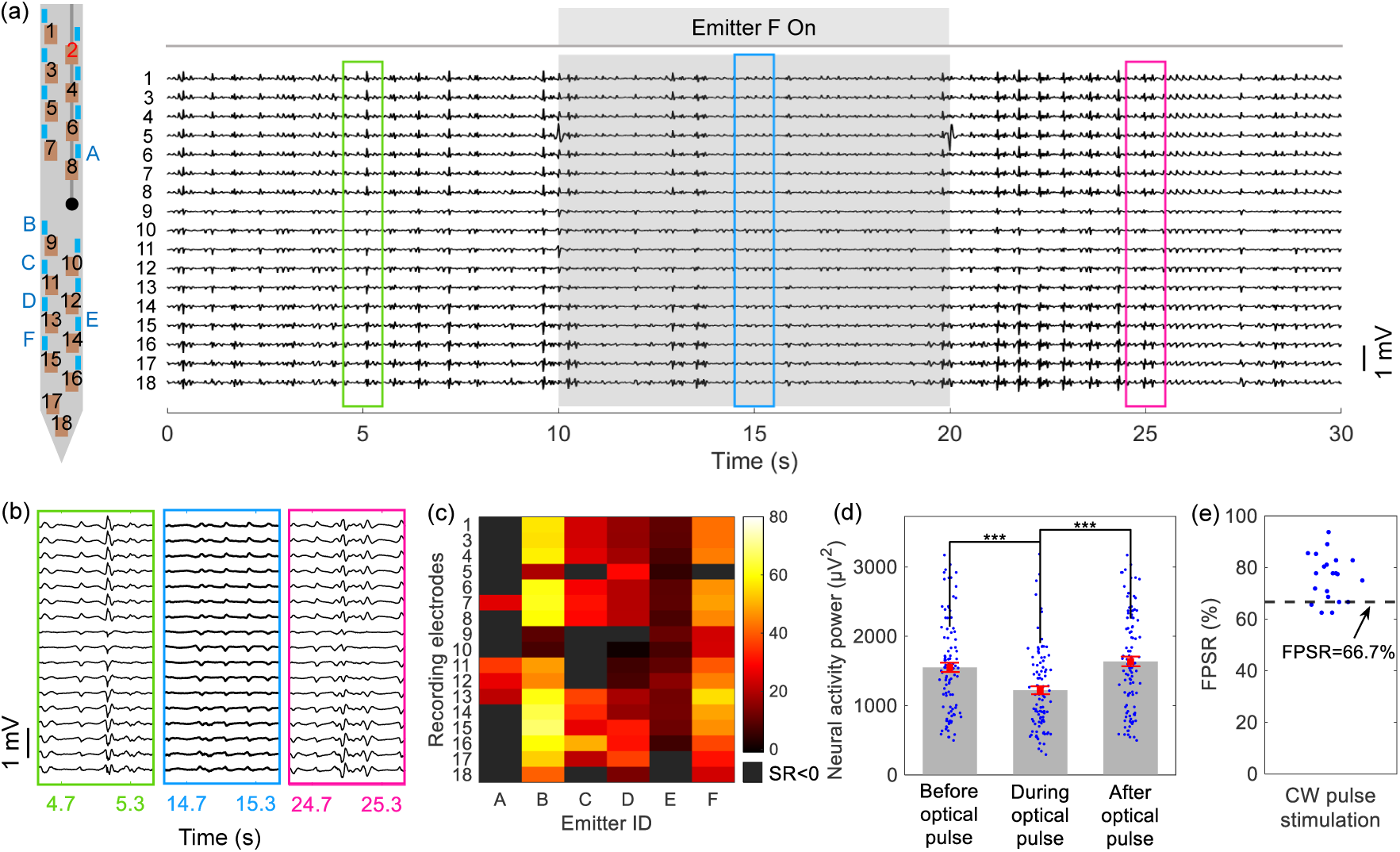
*In vivo* demonstration of full multifunctional neural probe operation, including seizure induction via microfluidic 4-AP injection, photostimulation for local seizure suppression, and electrophysiological monitoring. (a) (Left) Probe shank schematic with electrodes (1 - 18) and selected emitters (A - F) labeled; electrode 2 was found to be non-functional during the experiment. (Right) LFP traces from 17 electrodes following the onset of seizure. The recording window spans a 10-s continuous wave (CW) photostimulation pulse in addition to periods before and after, showing reduced LFP amplitudes with light delivery. Photostimulation parameters: Emitter F, ≈ 3.6 µW emission power, 10 s optical pulse. The displayed LFP traces were processed with common average referencing and bandpass filtering (5 - 300 Hz). (b) Magnified sections of the LFP traces in (a); before (green), during (blue), and after (magenta) photostimulation. (c) Heatmap of seizure suppression ratios (*SR*s) over a single photostimulation pattern; one pulse delivered in sequence from each of the selected emitters, recordings and *SR* calculations from each of the 17 electrodes. *SR >* 0 indicates suppressed seizure activity during the CW pulse. (d) Neural activity power calculated from LFP signals before, during, and after the optical pulse (n=102); each data point corresponds to an electrode and selected emitter. One-tailed non-parametric Mann–Whitney test was applied, with *** denoting statistical significance with p < 0.001. The error bars denote SDs. (e) Fractions of positive suppression ratios (*FPSRs*) during the CW photostimulation from all trials in the 5 animal experiments (n=21).

To evaluate the extent of seizure suppression in the LFP recordings, we define the suppression ratio (*SR*) of a seizure as the percent reduction in neural activity power (*P*) during photostimulation (see Eq. 1) [56], where *P_pre_* and *P_dur_* are the neural activity power before and during stimulation, respectively. The neural activity power of an LFP signal is defined in Eq. 2, where *v*(*t*) is LFP amplitude (sampled at times *t_n_*) and *T* is the number of time samples.

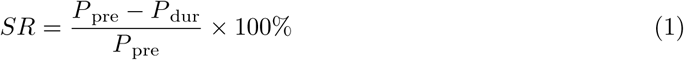

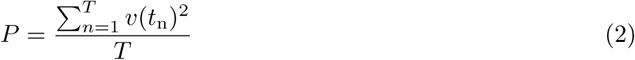

A positive *SR* corresponds to reduced LFP amplitude and/or frequency, indicating suppression of seizure activity, while a negative *SR* suggests an increase in seizure severity. Figure 6c summarizes the calculated suppression ratios over one instance of the photostimulation pattern as a heatmap. The 102 pixels of the heatmap correspond to 6 optical pulses (one from each of the selected emitters) with LFP signals recorded before and during each pulse (one signal from each of the 17 electrodes). 77.5% of these data points exhibited *SR >* 0 with a mean *SR* of 30%. Figure 6d compares the neural activity power from all recorded LFP traces before, during, and after each of the optical pulses of Fig. 6c. Reduced neural activity power during photostimulation is observed with statistical significance, while pre- and post-stimulation neural activity power are similar. The photostimulation patterns were repeated 3 times in the experiment presented in Fig. 6. Similar to Fig. 6c, the other 2 stimulation patterns resulted in 80.4% and 68.6% of the data points exhibiting *SR >* 0, with a mean *SR* of 22% and 40%, respectively.

Across these experiments, photostimulation from some emitters resulted in increased neural activity power on some electrodes. For example, in Fig. 6c, 22.5% of data points exhibited *SR <* 0. This observation aligns with reports of paradoxical effects of photostimulation in seizure suppression/promotion due to the interplay between interneurons (inhibitory) and pyramidal cells (excitatory) [51, 57–59]. To further quantify the effects of the photostimulation protocol used here, we define the fraction of positive suppression ratios (*FPSR*) as the percentage of positive *SR*s within one photostimulation pattern. *FPSR >* 50% indicates that the majority of LFP signals recorded during photostimulation (electrode-emitter combinations) exhibited suppressed seizure activity, while *FPSR <* 50% corresponds to seizure promotion across the majority of LFP signals. *FPSRs* of 21 photostimulation patterns across 5 *in vivo* experiments are shown in Fig. 6e. Among the trials, 18 displayed *FPSR* ≥ 66.7%, demonstrating repeatable local seizure suppression across ≥ 2*/*3 of emitter-electrode combinations.

Additional tests were performed to investigate the origin of the observed seizure suppression. First, in the above-mentioned experiments, photostimulation patterns applied prior to the first injection of 4-AP resulted in a broad increase in neural activity across 4 *in vivo* experiments, in contrast to the seizure suppression response to photostimulation observed following 4-AP injections (see Fig. S3). Second, separate experiments with wild-type mice demonstrated seizure induction with microfluidic 4-AP injections and no obvious change in neural activity power during photostimulation (see Fig. S4), excluding heating effects with light delivery as a cause of the observed light induced local seizure suppression. Third, following Refs. 56, 57, and 60, we tested low-frequency (5-ms pulses at 1 Hz, and 50-ms pulses at 5 Hz) and high-frequency (5-ms pulses at 20 Hz) photostimulation protocols. Suppression of seizure activity was not observed in these cases (see Fig. S5).

From the outset, given that (excitatory) pyramidal cells comprise the vast majority of ChR2+ cortical neurons in the Thy1-ChR2-EYFP mice used in this work [40], increasing neural activity is expected with photostimulation and, in the absence of seizure activity, this behavior was confirmed (see Section 2.2 and Fig. S3). With seizure activity, the observed inhibitory response to photostimulation opposes this expectation, and with the results of the above-mentioned additional tests, we posit the following hypotheses for this outcome. Our first hypothesis follows the lack of seizure suppression observed with low- and high-frequency pulse trains (consisting of pulses substantially shorter than the 10-s CW pulses used in Section 2.4) — suggesting long pulse durations as a critical factor to the observed seizure suppression. Firing rate fatigue [61] and depolarization block [62] are known mechanisms through which extended periods of photostimulation may lead to inhibition of excitatory cells. This is also supported by the results of our prior work on nanophotonic neural probes [26], where we observed inhibition following 1-s-long light pulses in Thy1-ChR2-EYFP mice. In this hypothesis, contributions to these inhibitory mechansisms from high baseline levels of activity during seizures is the distinguishing factor between excitation (inhibition) with photostimulation in the absence (presence) of seizure activity. Our second hypothesis (not mutually-exclusive to the first) is the secondary activation of interneurons following photostimulation of pyramidal cells during seizures resulting in an overall inhibitory response, with interneuron-mediated network activity being a known cause of paradoxical photostimulation effects [57, 63].

## 3 Discussion and Conclusion

In this work, we have demonstrated foundry-fabricated, multimodal, implantable, Si nanophotonic neural probes with photostimulation, electrophysiological recording, and microfluidic injection functionalities. In our integration approach, buried microfluidic channels are routed beneath dense electrical and photonic layers, enabling one microfluidic channel, 18 microelectrodes, and 16 grating coupler emitters on a single implantable shank. To the best of our knowledge, this configuration provides the highest number of addressable emitters reported to date in a neural probe integrating light delivery, electrophysiology, and microfluidics. The ≈1-mm span of the emitter–electrode array provides substantial coverage compatible with individual mouse brain structures (e.g., cortex or hippocampus), while the integrated microfluidic channel enables complementary drug delivery. In comparison with prior technologies (Table 1), Si-based multimodal probes with higher electrode counts have been demonstrated but were limited to a single polymer waveguide [13] or a single co-packaged fiber emitter [24]. Polymer-fiber approaches have supported up to 8 emitters [19] and 6 electrodes [12], but at the cost of cross-sectional areas ≈1.9–9.8× larger than in this work, leading to greater tissue displacement and damage. Further reductions in our probe dimensions are feasible, as we have demonstrated in Ref. 5, with 30-µm-thick shanks via backside polishing of singulated chips. Parallel to our monolithic microfluidic integration, we also reported in Ref. 64 a nanophotonic probe with a 3D-printed microfluidic channel atop the shank (not included in Table 1 as recording electrodes were not tested). Compared to the current work, these 3D-printed channels were restricted in outlet placement and required additional post-processing beyond foundry fabrication. Looking forward, the prototype neural probes demonstrated here establish a foundation for extending functionality (e.g., scaling photonic circuit complexity, reducing optical losses, and increasing the number of microfluidic channels) and enhancing usability in neuroscience experiments (e.g., through miniaturized packaging). In the following, we expand on these directions and highlight neuroscience use cases for both current prototypes and future realizations.

The number and density of emitters on the neural probe define the spatial resolution and coverage of photostimulation, with higher values enabling interrogation of larger neuronal populations at greater precision. The emitter density demonstrated here is a result of the sub-micron dimensions and high optical confinement of the SiN waveguides within our nanophotonic neural probes, supporting closely-spaced arrays of on-shank waveguides and compact grating coupler emitters. However, the number of emitters is limited by the number of cores in the multicore fiber, each coupling light to an on-chip waveguide. While increasing the number of fiber cores would directly enable addressing of more emitters, practical limits on the fiber diameter pose challenges for marked scaling of the emitter count. Alternatively, advanced passive devices [27, 29] and optical switches [3, 65] may be realized with nanophotonic SiN waveguides and the accompanying electrical layers, providing additional pathways for scaling the number of emitters. Through integrated wavelength demultiplexer devices, such as ring resonators [23] or evanescent couplers [5], multiple emitters may be addressed by each routing waveguide through control of the laser wavelength. Also, networks of thermally-actuated 1×2 optical switches may be used to spatially address arrays of grating coupler emitters [3, 4]. Each of these approaches enables multiplicative scaling of the number of addressable emitters per fiber-to-chip edge coupler and corresponding fiber core. Our Si photonics foundry process can support such scaling, as optical switches and advanced passive devices, some leveraging multiple waveguide layers, have recently been demonstrated in our visible-light photonics platform [5, 29, 65].

In optogenetics, maximum optical output powers of ∼100 µW–1 mW per emitter are desirable, from low levels for single-neuron stimulation near the shank to higher levels for network activation and extended illumination in scattering tissue [4, 25, 26]. Achieving these levels will require improvements in both on-chip photonic circuitry and packaging, as the neural probes reported here exhibited high optical loss (*>*25 dB), primarily due to scattering from edge couplers and polarizers. Post-packaging, the overall optical loss increased to ≈30 dB, a result of misalignment between the multicore fiber and the array of edge couplers during and/or after the packaging process, limiting emission powers to ≈2.4 - 6.4 µW. Variations between designed and fabricated waveguide thicknesses and widths are a likely cause of high on-chip losses, leading to optical mode mismatches between our inverse taper edge couplers and the multicore fiber in addition to increased bend losses in the polarizers. Reduced device losses are expected through improved control of waveguide dimensions in fabrication and the introduction of variation-tolerant designs, e.g., adiabatic polarizers as in Ref. 66. Meanwhile, improved fiber-to-chip alignment is anticipated through optimization of the attachment process to reduce epoxy shrinkage during UV curing. Further engineering at both the chip and packaging levels, together with the use of higher-power lasers, is expected to increase emission powers. As a step in this direction, in Ref. 5, we reported a dual-color nanophotonic probe (without microfluidics) achieving output powers *>*80 µW from individual emitters.

While the multimodal neural probe reported here features a single embedded microfluidic channel, the fabrication process (Fig. 2a) can, in principle, accommodate multiple input ports, channels, and outlets within one probe for multi-chemical delivery and sampling. In the current single-channel neural probe design, injecting a second chemical during *in vivo* experiments requires retraction of the neural probe, flushing of the channel, and loading of the new solution. This process typically takes over 30 minutes and necessitates a second insertion of the neural probe, leading to additional tissue damage and potential variability between injection locations. By contrast, probes with multiple microfluidic channels can enable rapid switching between different chemicals within a single implantation [17], simplifying experimental procedures, minimizing tissue damage, and allowing diverse combinations of drug delivery. Moreover, multi-channel neural probes can enable simultaneous chemical delivery and neurochemical sampling from distinct pathways within the same implant. In Ref. 16, one microfluidic channel delivers high-potassium artificial cerebrospinal fluid to stimulate neurochemical release, while another channel collects extracellular fluid for offline analysis with mass spectrometry to monitor resulting changes in neurotransmitter concentration (e.g., dopamine and *γ*-aminobutyric acid). We envision future generations of our multimodal nanophotonic neural probes integrating multiple microfluidic channels for precise manipulation and monitoring of neurochemical dynamics *in vivo*. Based on the cross-sectional image in Fig. 2f and assuming a minimum gap of 10 µm between adjacent channel edges, we estimate an on-shank minimum channel pitch of ≈22 µm, with the shank width of the current probe design potentially supporting 4 channels.

In the packaging process of the multimodal neural probes, optical fibers and fluidic tubing are permanently attached to the probe chips, necessitating close proximity to the peripheral control system and restricting animal movement during experiments. Toward flexibility in experiments with head-fixed animals and, potentially, long-term chronic experiments with freely moving animals, the introduction of multicore fiber connectors [67] is a possible means of realizing detachable optical interfaces. Further miniaturization via integration with on-chip laser diodes would eliminate the need for external laser sources and optical fibers [68] and toward this goal, we have recently demonstrated flip-chip integration of blue laser diodes within visible-light photonic integrated circuits [69]. In parallel, wireless microfluidic systems offer a compelling alternative to the restrictions imposed by fluidic tubing and external flow control systems. Miniaturized assemblies of refillable chemical reservoirs and microscale pumps (e.g., with thermal actuation [70]) present possibilities for future generations of wireless microfluidic neural probes. Combining these strategies for optical and fluidic packaging could enhance usability in experimental neuroscience while enabling long-term studies, including drug efficacy testing, repeated local delivery [70], and behavioral assays [15].

*In vivo* validation of our multimodal neural probes was performed in optogenetic mice sensitive to blue light, demonstrating neural activation with photostimulation, seizure induction via microfluidic drug delivery, and concurrent electrophysiological recordings. Simultaneous multimodal operation was further shown through seizure induction and local suppression using drug injections and photostimulation, respectively, as confirmed with LFP recordings. Beyond validating functionality, these demonstrations highlight potential use cases in epilepsy research. Neuroscience experiments studying the role of specific cell types in seizure generation/cessation [71] and mapping of seizure propagation pathways [59] stand to benefit from the cell-type-specific optogenetic stimulation, targeted and on-demand seizure induction, and spatially-resolved recording capabilities of multimodal nanophotonic neural probes. By contrast, conventional approaches that rely on separate drug injections and implanted Si probes, fibers, or electrodes [50, 51, 59] are limited by lower spatial accuracy, increased tissue displacement, and higher experimental complexity due to multiple implants. Additionally, the reported multimodal neural probes hold potential for a wider range of neuroscience applications, including optical uncaging of locally injected caged neurotransmitters (e.g., caged glutamate [72] and dopamine [73]); targeted viral opsin injection with subsequent co-localized optical stimulation and neural recordings [24]; validation of pharmacological therapeutic models; and integration with complementary benchtop or implantable neurotechnologies (e.g., for fluorescent functional imaging of seizure dynamics [74] or evaluation of electrical and ultrasound-based neuromodulation strategies [75]).

In their current form, the prototype multimodal neural probes reported here already support new experiments, though restricted to head-fixed configurations. Building on this work, these probes could offer multiple experimental degrees of freedom for advanced studies of seizure dynamics, including mapping propagation with the integrated microelectrode array, triggering seizures via optogenetic stimulation [26] or kainic acid administration [50] (leaving the probe’s microfluidic channel available for other agents), testing seizure-suppressing compounds via the microfluidic channel [46], and applying the platform across multiple optogenetic mouse strains to optically probe the roles of different cell types. Because the neural probes combine multiple functions on a single shank, experiments can also take advantage of available space to implant other complementary devices nearby or in different brain regions, as demonstrated in Ref. 5, where a high-density CMOS electrophysiology probe was implanted alongside a nanophotonic probe. With the addition of one extra microfluidic channel, push–pull fluid sampling may be achieved [16, 17], enabling a range of neurochemical sensing applications. Future implementations incorporating chip- and packaging-level innovations outlined above are expected to expand these capabilities further, critically enabling chronic implantation.

In conclusion, we have demonstrated multimodal nanophotonic neural probes with optogenetic stimulation, electrophysiological recording, and microfluidics capabilities. Our monolithic integration approach defines nanophotonic waveguides and metal wiring atop buried microfluidic channels, opening an avenue toward parallel layers of dense photonic, electrical, and fluidic routing. The prototype neural probes featured 16 addressable emitters, 18 microelectrodes, and one buried microfluidic channel on a single shank, setting a record for the number of emitters among neural probes combining light emission, recording, and microfluidics functionalities. Paths towards scaling the number of emitters, integrating multiple microfluidic channels, and miniaturizing the overall assembled systems were outlined. A series of *in vivo* experiments validated each modality individually and in combination, and applications to epilepsy research were highlighted. The neural probes were fabricated in a commercial Si photonics foundry, offering a direct route to scaling of manufacturing volumes and distribution to neuroscientists en masse. With continued development, we envision future generations of this neurotechnology offering higher optical, electrical, and fluidic channel counts in a smaller implant form factor, positioning these probes as versatile tools for neuroscience and translational research.

## 4 Materials and Methods

### 4.1 Electrode post-processing for impedance reduction

Similar to our previous work [5, 26], TiN microelectrode surfaces were post-processed (roughened) by femtosecond laser pulses [76] to achieve reduced electrochemical impedances (*<*2 MΩ) for neural recordings with high signal-to-noise ratios [39]. Each neural probe was sandwiched between a glass slide and a thin coverslip; the edges of the coverslip were taped for mounting and sealing. Drops of deionized water were added on top of the coverslip for imaging with a water-immersion objective (N16XLWD-PF, Nikon Instruments Inc., Melville, NY, USA) installed on a two-photon microscope (Ultima 2Pplus, Bruker, Billerica, MA, USA). The field of view (FOV) of the microscope scan pattern was adjusted such that only one electrode was included and centered in the FOV during each laser treatment. Femtosecond laser pulses from a 1035-nm-wavelength pulsed laser (Coherent Monaco 1035, ≈270 fs pulse width, 10-kHz repetition rate, averaged optical power at sample of ≈60 µW) were scanned across each TiN electrode surface. Following the post-processing procedure, each neural probe was inspected under an optical microscope, with sucessfully processed electrodes appearing darkened. Following packaging of the neural probes (Section 4.2), electrode impedances were measured at a 1 kHz frequency by dipping the shank of each neural probe in a 1× PBS solution together with a Ag/AgCL reference electrode; impedance measurements were acquired with an Open Ephys data acquisition system (Open Ephys Production Site, Lisbon, Portugal) via an electrophysiology headstage circuit board (mini-amp-64, Cambridge NeuroTech, Cambridge, UK).

### 4.2 Neural probe packaging

For each neural probe, the packaging process began with attaching a thin metal spacer block (approximately of equal width and length to the base of the probe chip) to a custom PCB (see Fig. 3a). The neural probe was then attached onto the metal spacer. Both attach steps were performed with thermally cured silver epoxy (Ablebond 84-1LMIT1, Loctite, Stamford, CT, USA). The metal spacer positioned the neural probe at an appropriate height (≈400 µm) above the PCB surface for attaching a multicore fiber to the neural probe. The PCB was pre-assembled with two electrical connectors (Molex SlimStack, .35mm SSB6 PLUG 34CKT) for connection to the headstage circuit board. Bond pads on the neural probe chip were wire-bonded with aluminum wire to corresponding pads on the PCB. The bonded wires were encapsulated with a UV-curable epoxy (Katiobond GE680, Delo, Germany).

Next, a 23-gauge, 90°-bent, stainless steel microtube (inner diameter ≈337 µm, outer diameter ≈641 µm, PN-BEN-23G-20, Darwin Microfluidics, Paris, France) attached to a length of silicone tubing (LVF-KTU-13, Elveflow, Elvesys, Paris, France) was aligned and coupled to the fluidic input port of the neural probe chip (see Fig. 2d). The position of the microtube was adjusted by a 5-axis piezoelectric alignment stage. With the fluidic input port of the neural probe positioned within the steel microtube, UV-curable epoxy (Katiobond GE680, DELO, Munich, Germany) was applied and cured at the interface between both, forming a sealed fluidic connection to the on-chip microfluidic channel.

Finally, a custom 16-core optical fiber [37] (mounted in a fiber ferrule) was aligned and attached to the array of 16 edge couplers at the neural probe chip facet using a 6-axis piezoelectric alignment stage and UV curable epoxies (OP-4-20632 and OP-67-LS, Dymax Corp., Torrington, CT, USA); the epoxies were applied and cured in sequence. The base of the neural probe chip, fiber ferrule, and wire bonds were subsequently encapsulated in optically opaque epoxy (EPO-TEK-320, Epoxy Technology, Billerica, MA, USA). Additional details of the fiber-to-chip attachment process are available in Ref. 26.

### 4.3 Neural probe system: electrophysiological recording and optical functions

Figure S1 illustrates the electrophysiological recording and optical subsystems of the overall neural probe system. In the recording subsystem, extracellular electrophysiological signals were routed to bond pads on the base of the neural probe chip, and wire bonds transmitted the signals to the PCB (on which the probe was attached). The PCB traces routed the signals to electrical connectors to which an electrophysiological recording headstage (mini-amp-64, Cambridge NeuroTech, Cambridge, UK) was connected. A co-axial electrical cable (C3203, RHD standard SPI interface cable, Intan Technologies, Los Angeles, CA, USA) routed amplified and digitized signals from the headstage to an Open Ephys acquisition system (set to capture wideband signals, 1 - 7500 Hz at a 30 kHz sampling rate per channel).

The optical subsystem enabled addressing of the emitters on the neural probes and definition of photostimulation patterns. Similar to our previous work in Refs. 26 and 31, the neural probe was connected to a computer-controlled laser scanning system, Fig. S1, through a multicore fiber. The laser scanning system directed and focused light from a 488-nm-wavelength diode laser (06MLD-488, Cobolt, Solna, Sweden) to the 16 cores of the multicore fiber via a microelectromechanical system (MEMS) mirror (A7B2.1-2000AL, Mirrorcle Technologies Inc., Richmond, CA, USA) in addition to free-space focusing and relay optics. Actuation of the MEMS mirror enabled selection of the fiber core to which light was coupled, and correspondingly, the on-chip waveguide and grating coupler guiding and emitting light. To prevent emission power fluctuations from the probe due to polarization fluctuations in the multicore fiber and polarization-dependent transmission of the on-chip photonic circuitry, laser light input to the scanning system was depolarized (using a free-space depolarizer [26]). The on-chip polarizers of the neural probe (see Section 2.1) selected the TE-component of the depolarized light, enabling a stable, on-chip polarization state. Additionally, the optical power input to the scanning system (and the emitted power from the probe) was controlled with a free-space variable attenuator mounted in a motorized rotation mount. An optical shutter formed the optical pulses of the photostimulation patterns, and digital modulation of the laser turned the emission on(off) during(between) photostimulation patterns.

A laboratory computer provided control of the experimental apparatus and logging of data. A custom software interface in MATLAB (MathWorks Inc, Natick, MA, USA) enabled programmatically defined photostimulation patterns and emitter addressing. The shutter and laser were computer controlled via a microcontroller (Teensy 3.6, PJRC, Sherwood, OR, USA), which generated transistor-transistor-logic (TTL) control signals. The microcontroller also transmitted photostimulation pulse timestamps to the Open Ephys system. Electrophysiological recordings were monitored and stored in the computer using the Open Ephys graphical user interface (GUI). Microfluidic injections were also controlled with the laboratory computer through communication with the flow controller (see Section 4.4).

### 4.4 Neural probe system: microfluidic injection

Figure S1 also illustrates the fluidic subsystem of the overall neural probe system. A pressure-driven flow controller (OB1 MK3+, Elveflow, Elvesys, France) with a pressure channel permitting working pressures up to 8 bar was used to precisely drive and control the microfluidic injections. A compressed air source up to 3.5 bar was used as the pressure source of the flow controller. A flow rate sensor (MFS2, Elveflow, Elvesys, Paris, France) was connected to the flow controller outlet with a silicone tube and fittings, enabling real-time flow rate data acquisition during the injections. The flow controller communicated with a laboratory computer featuring control software (Elveflow Smart Interface, Elveflow, Elvesys, Paris, France) for pressure/flow rate control and data acquisition. A feedback loop defined in the control software ensured flow regulation with stablized pressure and flow rate. On the day of the experiment, the 4-AP solution (in a 1.5-mL Eppendorf tube reservoir) was thawed, and the reservoir was connected to the outlet of the flow controller through an adapter and matching fittings (LVF-KPT-XS-2-2, Elveflow, Elvesys, Paris, France). The silicone tube attached to the neural probe was connected to the output port of the flow rate sensor. Overall, the flow controller regulated the pressure source and drove the solution from the reservoir through the in-line flow rate sensor, microfluidic channel of the neural probe, and into the brain. The pressure drop and flow rate in the fluid path were monitored and recorded in real time.

### 4.5 *In vivo* experimental procedure

All animal experiments were performed under protocols approved by the animal care committees of the University Health Network in accordance with the guidelines of the Canadian Council on Animal Care. Adult (2 to 4 months old) male and female Thy1-ChR2-EYFP (n=15, strain number 007612, The Jackson Laboratory, Bar Harbor, Maine, USA) and wild-type (n=2, C57BL/6J, Charles River Laboratories, Wilmington, Massachusetts, USA) mice were kept in a vivarium at 22°C (12-hour light/dark cycle, food and water available *ad libitum*).

1 - 7 days prior to the experiment, a Thy1-ChR2-EYFP or wild-type mouse was anesthetized (by induction with 5% and maintenance with 1% to 2% isoflurane/oxygen anesthetic) and mounted in a stereotaxic frame with ear bars (Model 902, David Kopf Instrument, Tujunga, California). Prior to skin incision, the surgical site was treated in sequence with betadine scrub, 75% alcohol, and betadine solution; preoperative analgesia was administered at the surgical area with a lidocaine injection (3% in saline, ≈0.1 mL) and buprenorphine (0.1 mg/kg, 0.05 - 0.1 mL per subcutaneous injection). Following skin incision, two skull screws for external reference/ground electrodes (19010-10, Fine Science Tools, Foster City, CA, USA) and a headplate were fixed to the mouse skull by 1) drilling two small holes in the skull with a dental drill, (2) inserting the screws in the holes, and (3) attaching the headplate to the skull with dental cement (C&B Metabond, Parkell, Edgewood, NY, USA). The location of probe insertion (AP: −0.5 mm, ML: 1.2 mm, targeting motor cortex) was determined with the sterotaxic frame and marked. The ground screw was mounted toward the frontal bone on the ipsilateral side of the probe insertion site, and the reference screw was toward the cerebellum on the contralateral side [26]. The mouse was placed back in the cage for post-operative recovery.

On the day of experiment, the mouse was first anesthetized (following the above procedure) and then transferred to the stereotaxic frame. Thin stainless steel wires were wrapped around the exposed threads of the reference/ground skull screws for connection with the headstage. Centered at the previously marked location for probe implantation, a craniotomy with a ≈2-mm diameter was performed. Dura was carefully removed for smooth insertion of the neural probe. Anesthesic was supplied throughout the surgery, and the mouse was transferred to a head-fixed measurement apparatus immediately after the surgical preparation. Before recovery from anesthesia, the mouse was placed in a 3D-printed cylinder (to restrict body movements) and the headplate was attached to metal bars (all within the measurement apparatus). The bone screws were attached to the ground and reference leads of the headstage.

The measurement apparatus included an upright epifluorescence microscope (MM201, Thorlabs Inc.) positioned above the mouse for imaging the probe insertion process. A packaged neural probe was mounted on a 4-axis motorized micromanipulator (QUAD, Sutter Instrument Company, Novato, CA, USA) for precise insertion at an angle ≈23*^◦^* from vertical. After positioning the shank tip in close proximity to the target implantation site, the neural probe was inserted into the brain (along the diagonal axis of the micromanipulator) at a speed of ≈1 mm/s over ≈500 µm for fast penetration. Following a ≈20 min recovery period, the probe was advanced at a lower speed of ≈1 µm/s until spontaneous activity was detected on multiple electrode channels. Spontaneous activity levels were monitored over an observation period of ≈20 min. Photostimulation and/or microfluidic injection was performed when the baseline activity remained stable over the observation period. In cases where marked reductions in baseline activity were observed, the neural probe was advanced to a larger depth where spontaneous activity was observed again and the observation period was repeated.

After the experiment, the neural probe was retracted and the shank was immersed in a 1% w/v Tergazyme (Z273287, Sigma-Aldrich, St. Louis, MO, USA) solution for ≈2 hours, followed by rinsing in a deionized water bath for ≈10 min. The animal was put under general anesthesia with 5% isoflurane, and after it was fully unconscious, it was transferred to a CO_2_ chamber with a flow rate of 30 - 70% for euthanasia. Following *in vivo* experiments with microfluidic injections (Sections 2.3 and 2.4), the mouse brain was extracted and 300 µm thick brain slices were prepared with a vibratome (Leica VT1200 S, Leica Mikrosysteme Vertrieb GmbH, Germany). Epifluorescence imaging was performed to identify the insertion tract and confirm that microfluidic injections were successfully conducted [via red fluorescence of the Alexa Fluor 568 conjugated dextran dye (D22912, Invitrogen) in the injected solution]. Separate images of YFP and Alexa Fluor 568 dye fluorescence were acquired.

### 4.6 Beam profile measurement in brain slices

Emitted beam profiles from a packaged neural probe were measured in a brain slice via imaging of tissue fluorescence excited by the beams, Fig. 3d. A 1.5-mm-thick fresh brain slice was prepared (using a vibratome) from a Thy1-ChR2-EYFP mouse (4 months old). The selected brain slice (containing motor cortex) was attached to a glass slide with super glue (Insta-cure+, Bob Smith Industries, Atascadero, CA, USA). Drops of artificial cerebrospinal fluid solution were placed on and around the brain slice to prevent drying of the tissue. For imaging, the brain slice was mounted under an epifluorescence microscope with a 10× objective (M Plan Apo 10×, NA=0.28, Mitutoyo Deutschland GmbH, Germany), a green fluorescent protein (GFP) filter cube, and a CMOS camera. The neural probe was mounted on a 4-axis micromanipulator and inserted into the brain slice at an angle ≈23*^◦^* from vertical (such that the emitted beam was parallel to the brain slice surface). Three insertion sites were selected to confirm the beam profile characteristics in the presence of spatially-varying optical scattering in the slice. The insertion depth of the probe was selected for a shallow (*<*100 µm) depth of the emitted beam below the tissue surface. Separate sets of fluorescence images were acquired with epi-illumination from the microscope [showing the shank position, Fig. 3d(left)] and without epi-illumination [showing only the emitted beam from probe, Fig. 3d(right)].

### 4.7 Electrophysiological data analysis

The electrophysiological data analysis was performed with custom scripts in Python (version 3.11) and MATLAB. For electrophysiological data in Section 2.2 and Fig. S2 (without 4-AP injections and seizure activity), the raw electrophysiological data was firstly processed with (1) common average referencing by subtracting the mean of signals from all recording electrodes within the brain; (2) bandpass filtering with a 300 - 6000 Hz filter; and (3) photostimulation artifact removal by blanking out 5 ms at the beginning and end of the optical stimulus. Spike sorting of the processed data was then completed with the SpyKING CIRCUS toolbox [77]. Manual curation of the automatically sorted spikes was conducted in the phy GUI [78] based on the spike amplitude (≥30 µV), spike count (≥200), and inter-spike interval ratio (≤0.5 for a refractory period of 2 ms) [79]. Units with noisy waveforms were classified as noise and other units that failed to meet the aforementioned quality metrics were classified as multi-unit activity. Splitting and merging operations were performed for the remaining units based on their similarity, waveforms, and temporal firing rate distribution. Only sorted single units satisfying all the quality metrics were included in further data analysis.

For processing of local field potential signals (Sections 2.3 and 2.4, Figs. S3 - S5), common average referencing and photostimulation artifact removal were performed as stated above, while the frequency range of the applied bandpass filter was modified to 5 - 300 Hz. Spike sorting was not performed on LFP signals. The neural activity power [56] calculated from the LFP signal was used to quantify local seizure suppression.

All statistical tests between different data groups were performed with non-parametric Mann-Whitney-U-tests unless otherwise stated. Statistical significance was reported for p*<*0.05.

## Code and Data Availability

Electrophysiological data reported in this study and codes for data analysis are available upon reasonable request from the corresponding authors.

## Acknowledgments

This work was supported by the Max Planck Society. The authors thank Dr. Liang Zhang at the Krembil Brain Institute for helpful discussions.

## Author Contributions

W.D.S conceived the initial concept for the neural probe technology. X.M. designed the neural probes with inputs from J.K.S.P and W.D.S. The fabrication process flow was designed by H.C., X.L., and W.D.S. H.C., X.L., and G.Q.L. were responsible for the fabrication of the wafers. X.M., F.C., J.N.S., and P.K. developed the probe assembly approach. H.M.C. conceived the application of the probes for epilepsy research. X.M., H.M.C., J.K.S.P., and W.D.S. designed the experiments. X.M., F.C., H.W., A.S., and W.D.S. built the experimental setups. M.M. and H.M.C. prepared the animal experiments. X.M. conducted the experiments and analyzed the results with guidance from F.C., H.M.C., and W.D.S. X.M. and W.D.S. co-wrote the manuscript with inputs from other co-authors. G.Q.L, J.K.S.P., T.A.V., and W.D.S. supervised the project.

## Supplementary Materials

### Multifunctional nanophotonic neural probe system

**Fig. S1:**
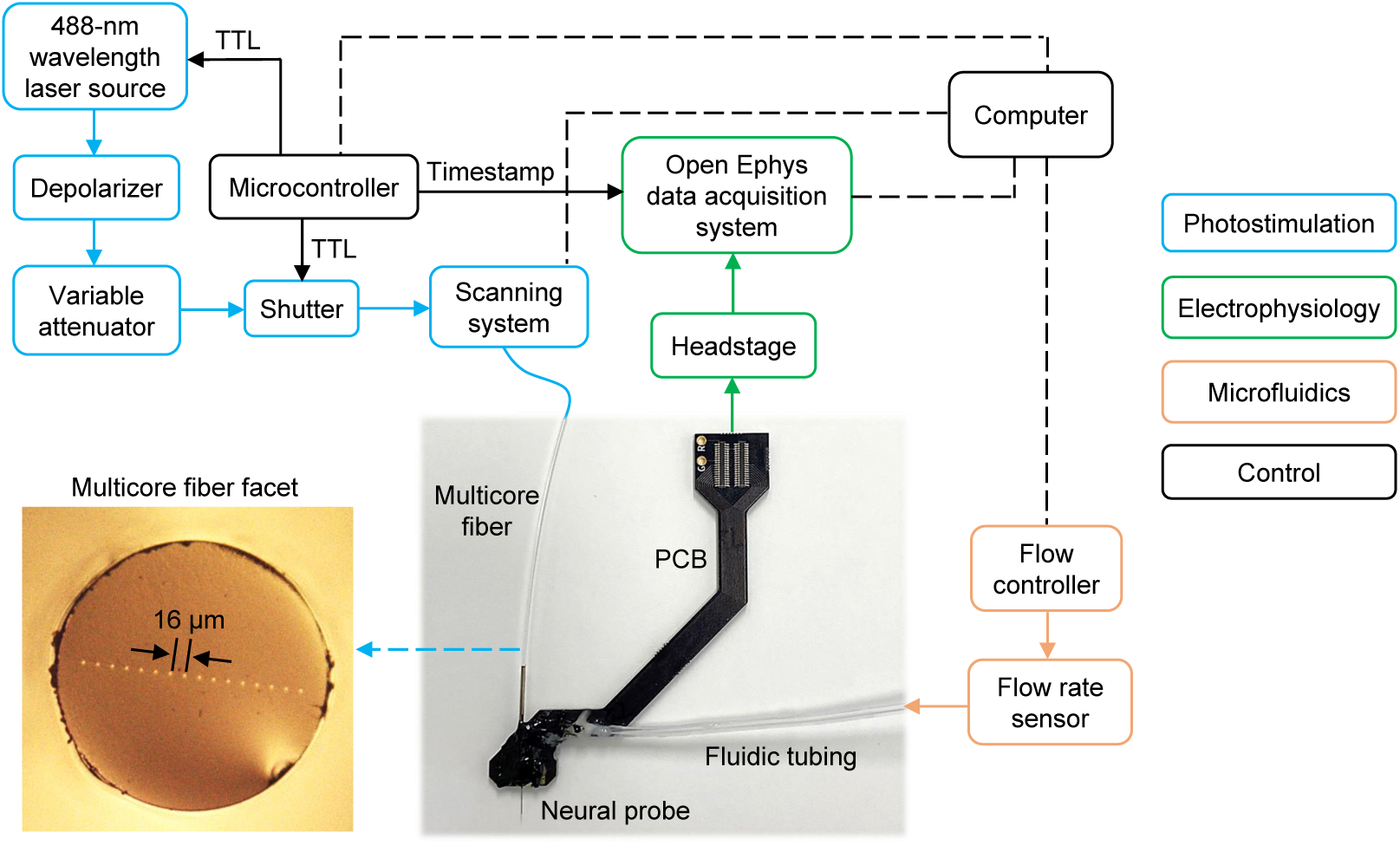
Schematic of the neural probe system. Laser light is transmitted from a 488-nm wavelength laser source to the neural probe via a depolarizer, variable optical attenuator, optical shutter, optical scanning system, and multicore fiber. The 16-core fiber is aligned and attached to the array of on-chip edge couplers of the neural probe. The scanning system enables selection of the fiber core (and optical emitter) to which light is coupled. The electrophysiological signals recorded by microelectrodes are acquired by an Open Ephys data acquisition system via a heastage and an electrical cable. A microcontroller controls the laser source and the optical shutter with transistor-transistor-logic (TTL) signals, defining the pulse trains for photostimulation. The TTL signals are also transmitted to the Open Ephys system to log the photostimulation timestamps. Fluidic delivery is actuated by a flow controller, with the in-line flow rate sensor capturing flow rates in real time. A laboratory computer provides overall system control through signals sent to the microcontroller, scanning system, and flow controller, while also retrieving recording and timing data from the Open Ephys system. Bottom-left: micrograph of multicore fiber facet. Bottom-middle: photograph of a packaged neural probe.

### Control experiments

**Fig. S2:**
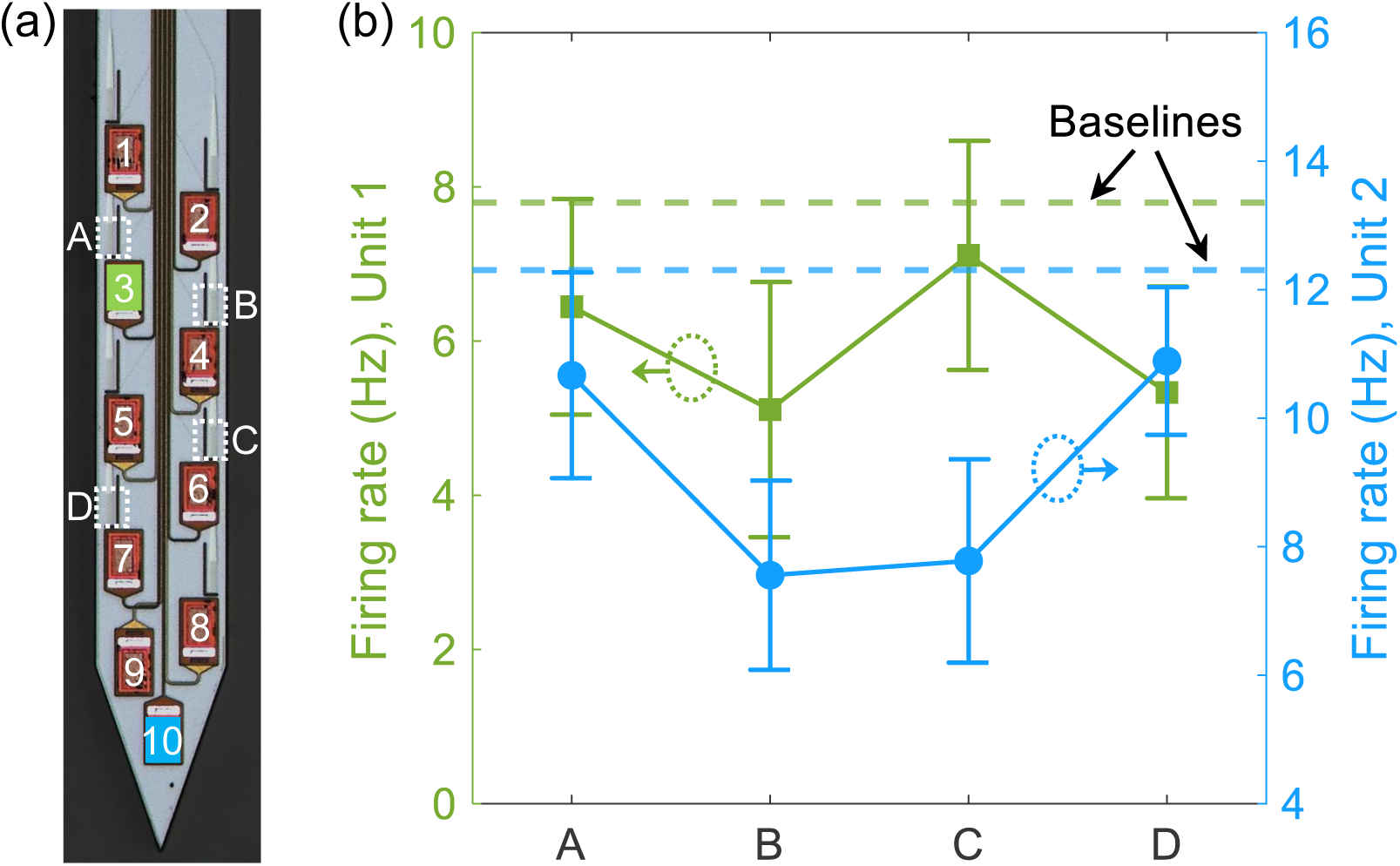
Photostimulation test in a wild-type mouse. (a) Micrograph of the neural probe shank with selected electrodes and emitters labeled. (b) Mean firing rates of 2 sorted single units (corresponding to electrodes 3 and 10) with photostimulation from 4 emitters (A - D). Mean baseline firing rates were averaged over 10 s before each stimulus. Error bars: standard error of the mean (SEM).

To verify the origin of the evoked spiking response with photostimulation from the neural probe in Section 2.2 (with blue-light-sensitive optogenetic mice), additional photostimulation tests were performed in two wild-type mice (2 to 4 months old). The implant location and photostimulation pattern were identical to Section 2.2, with the exception that a different set of four emitters were selected for sequential addressing (as shown in Fig. S2a). Also similar to Section 2.2, the emission powers ranged from ≈1.0 - 2.4 µW across the four emitters.

Figure S2b shows representative results from one of the control experiments, and similar results were observed in the other control experiment. Mean firing rates of two sorted single units (Unit 1 and Unit 2) with photostimulation from the four emitters are shown. Units 1 and 2 were detected on electrodes 3 and 10, respectively. Compared with the baseline firing rate, increases in firing rates during photostimulation were not observed, indicating the photostimulation effects reported in Section 2.2 were of optogenetic origin rather than a result of tissue heating with illumination. Five additional units were sorted in the experiment, and no significant changes in firing rates with photostimulation were observed.

**Fig. S3:**
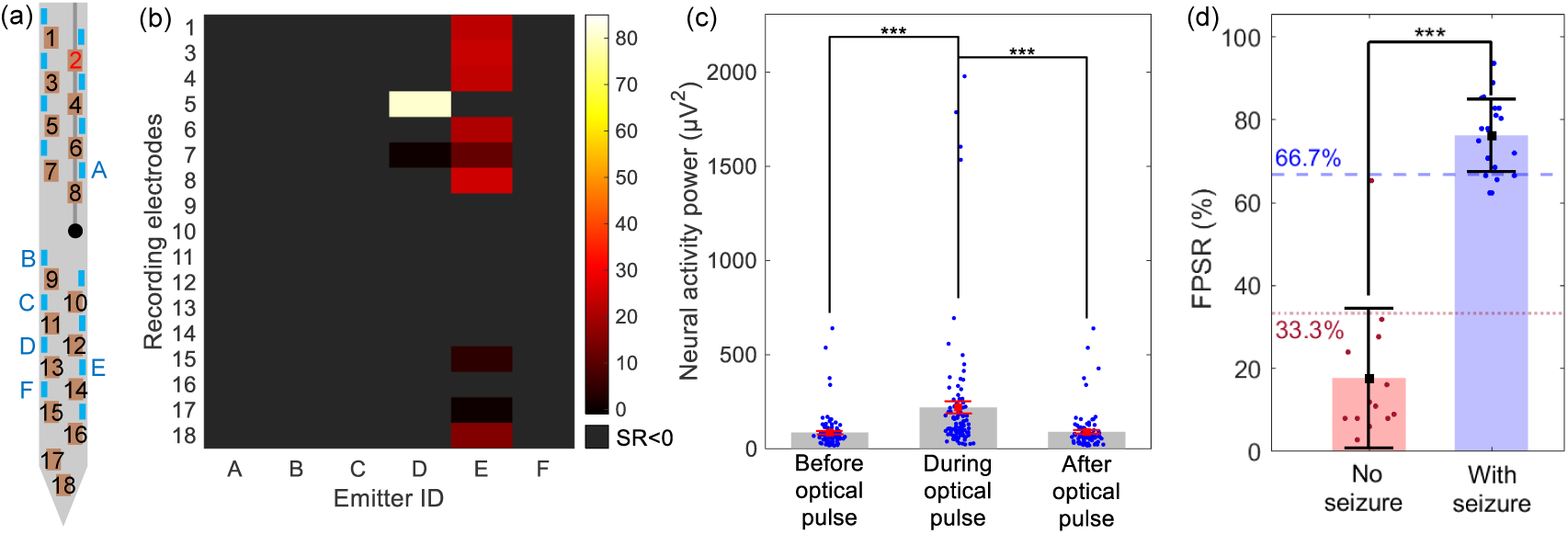
Control photostimulation tests prior to microfluidic 4-AP injection. (a) Schematic of the probe shank with electrodes and selected emitters labeled. (b) Heatmap of suppression ratios (*SR*s) on 17 electrode channels with CW optical pulses from 6 emitters. (c) Neural activity power calculated from LFP signals before, during, and after the optical pulses (n=102). (d) Fraction of positive suppression ratios (*FPSR*) in each photostimulation pattern in control tests (before 4-AP injections, n=13, 4 mice) and in seizure suppression tests (after 4-AP seizure induction, n=21, 5 mice, repeated from Fig. 6e). The error bars denote standard deviations in panels (c) and (d). One-tailed non-parametric Mann–Whitney test was used in panels (c) and (d). *** denotes statistical significance with p<0.001.

In four of the five *in vivo* experiments reported in Section 2.4, the photostimulation pattern was applied repeatedly before the first 4-AP injection. These control tests aimed to confirm the effects of photostimulation prior to the induction of seizure activity. Figure S3 shows the results from one of the four experiments (performed as part of the overall experiment in Fig. 6). The six selected emitters are shown in Fig. S3a and are identical to Fig. 6; the same emission powers were used. The heatmap of suppression ratios (*SR*, defined in Section 2.4) during the photostimulation pattern is shown in Fig. S3b. 91 of the data points (emitter-electrode combinations) exhibited *SR <* 0, indicating the photostimulation generally excited neural activity prior to the injections of 4-AP. Figure S3c compares the neural activity power before, during, and after the optical pulses. The neural activity power during photostimulation was significantly higher than pre- and post-stimulation, agreeing with the results of Section 2.2 and the expected photostimulation mechanism (excitation of Channelrhodopsin-2-positive pyramidal cells). As in Fig. 6d, neural activity power pre- and post-stimulation were similar. Also, due to the absence of seizure activity, the amplitude of the neural activity power was lower in these control tests compared to Fig. 6d.

Figure S3d summarizes the fraction of positive suppression ratios (*FPSR*, defined in Section 2.4) within each photostimulation pattern (trial). The set of control tests (13 trials with 4 mice) is compared to the set of seizure suppression tests in Section 2.4 (21 trials with 5 mice). During seizure suppression tests (following injections of 4-AP), 18 out of 21 trials had *FPSR* ≥ 66.7%. By contrast, in the control tests, 12 out of 13 trials exhibited *FPSR <* 33.3%, further indicating a general increase in neural activity with photostimulation prior to 4-AP injections.

**Fig. S4:**
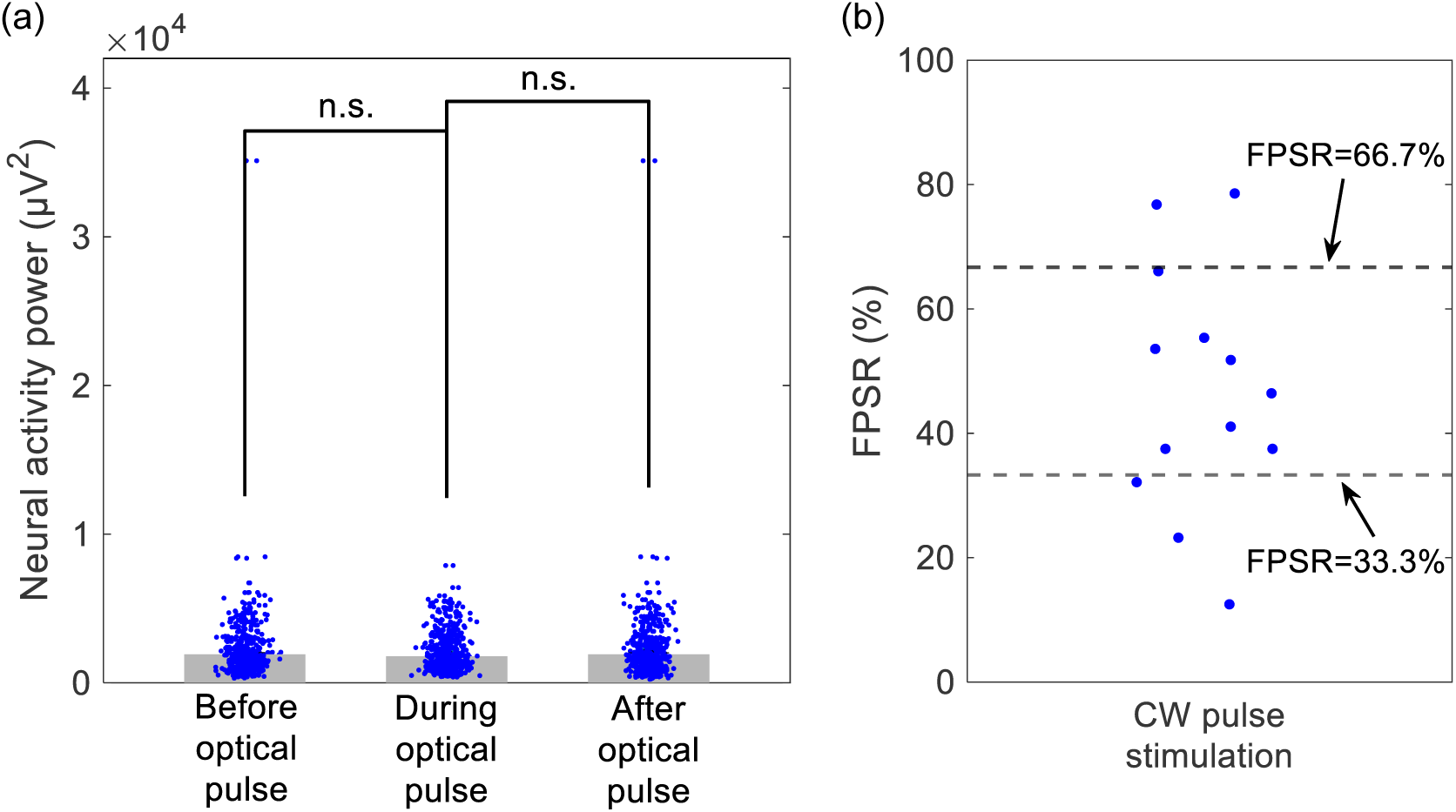
Seizure suppression tests in wild-type mice. (a) Neural activity power before, during, and after the 10-s continuous-wave optical pulses (n=392) in one mouse. The neural activity power was recorded from 14 electrodes during 7 photostimulation trials; light was applied sequentially from 4 emitters during each trial. Two-tailed non-parametric Mann–Whitney test was applied with n.s. denoting no statistical significance with p*>*0.05. (b) *FPSRs* calculated across 13 photostimulation trials in 2 wild-type mice. The error bars denote SEM.

Control experiments corresponding to the seizure suppression tests in Section 2.4 were performed on two wild-type mice (2 - 4 months old). The neural probe implant location, composition of the injected solution, and photostimulation pattern (10-s CW pulses emitted in sequence from each of the selected emitters) followed Section 2.4. Relative to Section 2.4, a different neural probe was used for these tests, and the set of selected emitters differed in location and number. Emission powers ranged from ≈1.0 - 4.2 µW. The neural probe captured LFP signals during seizure activity induced by the microfluidic injections of 4-AP, and Fig. S4 summarizes the recordings before, during, and after each photostimulation pulse. Seven repetitions (trials) of the photostimulation pattern were performed in the first wild-type mouse, and six were performed in the second mouse. At the implant depth of these experiments, 14 electrodes on the neural probe were within the brain. In Fig. S4a, no statistically significant difference was observed in the neural activity power before, during, and after the photostimulation pulses. Figure S4b shows the calculated *FPSRs* (defined in Section 2.4) across all photostimulation trials in both wild-type mice. 61.5% of the trials resulted in 33.3% *< FPSR <* 66.7%, indicating neither clear suppression nor promotion effect of seizure activity. Moreover, with *FPSR* ≥ 66.7% corresponding to clear suppression of seizure activity, only 15.4% of the control trials fell in this range, in contrast to 85.7% with the blue-light sensitive optogenetic mice in Section 2.4 — indicating that the local suppression of 4-AP induced seizure with CW optical pulses in Section 2.4 resulted from an optogenetic response instead of tissue heating with photostimulation.

### Tests of low- and high-frequency photostimulation patterns for local seizure suppression

**Fig. S5:**
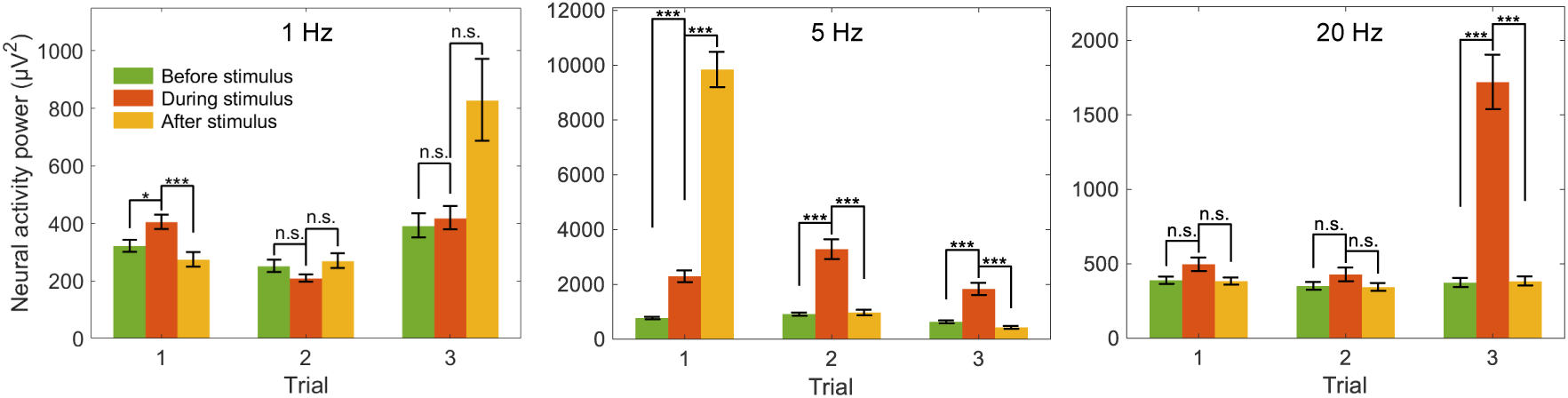
Neural activity power before, during and after photostimulation pulses at 1 Hz (left), 5 Hz (middle), and 20 Hz (right). Each photostimulation pattern was tested 3 times for each selected emitters during the experiment (trials 1 - 3). The error bars denote SEM. Two-tailed non-parametric Mann–Whitney test was applied with *** denoting statistical significance with p<0.001 and * for p<0.05, n.s. stands for no statistical significance with p≥0.05.

In addition to the CW photostimulation pattern tested in Section 2.4, additional photostimulation patterns from neural probes were tested for seizure suppression following microfluidic injections of 4-AP (with the same procedure of Section 2.4). These tests spanned low- (1 Hz and 5 Hz) and high- (20 Hz) frequency photostimulation pulse trains. The parameters of the three additional photostimulation patterns are summarized in Table S1; three different probes were used across the tests, and the number of selected emitters also varied. Three repeats of the pattern were performed for each selected emitter (trials 1 - 3). Figure S5 compares the neural activity power before, during, and after each stimulation pattern (calculated across all electrodes, as in Section 2.4). All tests were performed during seizure activity. Overall, no reduction in neural activity power during photostimulation (i.e. seizure suppression) was observed across the three additional photostimulation patterns.

**Table S1:** Summary of low- and high-frequency photostimulation patterns (adapted from the references listed)

| Frequency (Hz) | Pulse width (ms) | Recovery time (s) | Number of pulses | Number of emitters | Emission power range ($\mu W$) | Ref. |
| --- | --- | --- | --- | --- | --- | --- |
| 1 | 5 | 120 | 120 | 4 | $\approx 4.3 - 5.3$ | [57] |
| 5 | 50 | 120 | 600 | 1 | $\approx 3$ | [60] |
| 20 | 5 | 5 | 200 | 1 | $\approx 7.8$ | [56] |

